# Induced ligno-suberin vascular coating and tyramine-derived hydroxycinnamic acid amides restrict *Ralstonia solanacearum* colonization in resistant tomato roots

**DOI:** 10.1101/2021.06.15.448549

**Authors:** Anurag Kashyap, Montserrat Capellades, Weiqi Zhang, Sumithra Srinivasan, Anna Laromaine, Olga Serra, Mercè Figueras, Jorge Rencoret, Ana Gutiérrez, Marc Valls, Nuria S. Coll

## Abstract

The soil borne pathogen *Ralstonia solanacearum* is the causing agent of bacterial wilt, a devastating disease affecting major agricultural crops. *R. solanacearum* enters plants through the roots and reaches the vasculature, causing rapid wilting. We recently showed that tomato varieties resistant to bacterial wilt restrict bacterial movement in the plant. In the present work we go a step forward by identifying the physico-chemical nature of the barriers induced in resistant tomato roots in response to *R. solanacearum*. We describe that resistant tomato specifically responds to infection by assembling *de novo* a structural barrier at the vasculature formed by a ligno-suberin coating and tyramine-derived hydroxycinnamic acid amides (HCAAs). On the contrary, susceptible tomato does not form these reinforcements in response to the pathogen but instead displays lignin structural changes compatible with its degradation. Further, we show that overexpressing genes of the ligno-suberin pathway in a commercial susceptible variety of tomato restricts *R. solanacearum* movement inside the plant and slows disease progression, enhancing resistance to the pathogen. We thus propose that the induced barrier in resistant plants does not only restrict the movement of the pathogen, but may also prevent cell wall degradation by the pathogen and confer anti-microbial properties.

## Introduction

In natural environments plants are constantly exposed to diverse microbiota, including pathogenic organisms. In addition to pre-existing structural cell barriers that act as a first line of defense (Serrano *et al*., 2014; Falter *et al*., 2015), pathogen perception results in activation of a complex, multi-layered immune system in plants (Jones and Dangl, 2006). As part of the suite of inducible defenses, *de novo* formation of physico-chemical barriers prevents pathogen colonization and spread inside the plant. Despite their importance, the exact composition of these barriers, as well as the mechanisms that lead to their formation in the plant upon pathogen invasion remain largely unknown.

The interaction between the soil-borne bacterial wilt pathogen *Ralstonia solanacearum* and tomato offers a paradigmatic scenario to study inducible physico-chemical barriers, because of its agro-economic impact, and the well-developed genetic and molecular tools available in both organisms. *R. solanacearum* enters the root system through wounds or at the points of emergence of lateral roots, where the epidermal barrier may be compromised, and both the endodermis and Casparian strip are either not fully differentiated or reoriented and endodermal suberin overlying the primordium is being degraded (Vasse *et al*., 1995; Álvarez *et al*., 2010; Ursache *et al*., 2021) After entering the root, the bacterium moves centripetally towards the vasculature and once it reaches the xylem, it multiplies and spreads vertically within the vessels and horizontally to other vessels and the surrounding tissues (Digonnet *et al*., 2012).

The xylem tissue is in fact a major battleground for the interaction between vascular wilt pathogens and their hosts, where the outcome of the infection is at stake (Yadeta and Thomma, 2013). To prevent the spread of pathogenic propagules, the xylem vasculature of resistant plants undergoes intense structural and metabolic modifications. Resistant plants form vertical barriers such as tyloses and gels inside the vessel lumen, which in some plant-pathogen interactions effectively slow down vertical progression of the pathogen, or even confine it to the infection site, preventing systemic infection (VanderMolen *et al*., 1987; Rioux *et al*., 2018). Further, resistant plants also reinforce the walls of xylem vessels, pit membranes and surrounding xylem parenchyma cells in response to pathogens (Street *et al*., 1986; Benhamou, 1995). This prevents pathogen colonization of the surrounding parenchyma cells, nearby vessels and inter-cellular spaces through degeneration of the vessel pit membranes or cell walls (Nakaho *et al*., 2000; Digonnet *et al*., 2012) caused by the pathogeńs cell wall degrading enzymes (Liu *et al*., 2005; Pérez-Donoso *et al*., 2010; Lowe-Power *et al*., 2018). In addition, deposits in the xylem cell walls act as a shield against pathogen-derived metabolites such as toxins and enzymes, and diminishes water and nutrient availability for pathogens, thereby impeding their growth (Araujo *et al*., 2014). This vascular confinement is an effective strategy commonly found among plants resistant to vascular wilt pathogens such as *R. solanacearum,* which otherwise spread systemically once they reach the vasculature, clogging the vessels and causing irreversible damage and plant death (Potter *et al*., 2011; Scortichini, 2020; Kashyap *et al*., 2021;)

Among the various tomato germplasms, the cultivar Hawaii 7996 (H7996) is the most effective natural source of resistance against *R. solanacearum* (Nakaho *et al*., 2004; Grimault *et al*., 1994). In this cultivar, resistance to *R. solanacearum* is a complex polygenic trait. So far, two major (Bwr-12 and Bwr-6) and three minor (Bwr-3, Bwr-4, and Bwr-8) quantitative trait loci (QTLs) have been identified, although they only account for a portion of the observed phenotypic variation in resistance (2000; Thoquet *et al*., 1996; Mangin *et al*., 1999; Wang *et al*., 2013). Our previous study using luminescent and fluorescent reporter strains of *R. solanacearum* identified four distinct spatiotemporal bottlenecks through which H7996 is able to limit bacterial spread *in planta* (Planas-Marquès *et al*., 2019). In this resistant variety the pathogen encounters severe restriction in: i) root colonization, ii) vertical movement from roots to shoots, iii) circular invasion of the vascular bundle and iv) radial apoplastic spread from the vessels into the cortex. Vascular cell wall reinforcements seem to play a key role in confining *R. solanacearum* into the xylem vascular bundles of resistant tomato H7996. Ultra-microscopic studies in quantitatively resistant tomato cultivars showed that the pit membranes, as well as xylem vessel walls and parenchyma cells form a conspicuously thick coating in the form of an electron dense amorphous layer, as part of the defense response against *R. solanacearum* (Nakaho *et al*., 2000; Kim *et al*., 2016;). Thus, resistant H7996 plants have the ability to effectively compartmentalize *R. solanacearum* into the lumen of xylem vascular bundles. However, the type of barriers and compounds involved in this interaction remain understudied.

Among the polymers constituting vascular coating structures, lignin is the most typically found, constituting an integral part of the secondary cell wall of the xylem vasculature. Lignin has been well studied as a common structural defense against vascular wilt pathogens (Novo *et al*., 2017; Kashyap *et al*., 2021). Suberin has also been reported to be deposited in vascular coatings as a defense response (Kashyap *et al*., 2021), although the mechanisms regulating its synthesis, spatio-temporal dynamics and inducibility remain elusive. Interestingly, root microbiota has been recently shown to have the ability of shaping suberin deposits in the plant, highlighting its central role in plant-microbe interactions (Salas-González *et al*., 2021). Suberin is a poly(acylglycerol)-derived polyester containing long and very long chain fatty acid compounds and derivatives and also some aromatics, mainly ferulic acid, which is a hydroxycinnamic acid. Cells that accumulate suberin also accumulate lignin, whose deposition has been described to precede that of suberin in phellem cells (Lulai and Corsini, 1998). This lignin is also known as a lignin-like polymer. The lignin-like polymer consists of hydroxycinnamates and monolignols linked by C-C and ether bounds (Graça, 2015). The ligno-suberin heteropolymer formed by the lignin-like polymer and suberin has been also referred to as the poly(aromatic) and poly(aliphatic) domains of suberin, respectively. Commonly, suberized cell walls also comprise free fatty acyl derived compounds, known as suberin-associated waxes, and phenolic soluble compounds, which share biosynthetic pathways with suberin and lignin, respectively (Bernards, 2002).

Ferulic acid present in the suberin and lignin-like fractions is proposed to link both polymers (Graça, 2010) and its continuous production has been demonstrated essential for suberin deposition (Andersen *et al*., 2021). Ferulic acid amides, such as feruloyltyramine and feruloyloctopamine, have been described as structural components of the lignin-like polymer and in the phenolic soluble fraction of suberizing wound-healing potato tuber (Negrel *et al*., 1996; Razem and Bernards, 2002). Ferulic acid amides belong to the Hydroxycinnamic acid amide (HCAA) family, which present antimicrobial activity and are considered biomarkers during plant-pathogen interactions (Zeiss *et al*., 2021). However, the molecular, biochemical and physiological role of HCAAs in plant defense remains to be elucidated (Macoy *et al*., 2015). Besides their direct antimicrobial activity as soluble phenols, HCAAs have also been proposed to cross-link to cell wall structural polymers during infection, potentially contributing towards the formation of a phenolic barrier that can make the cell wall resilient to pathogenic degradation (Zeiss *et al*., 2021).

In the present study, we conducted a detailed investigation of the inducibility, structure and composition of the xylem vascular wall reinforcements that restrict *R. solanacerum* colonization in resistant tomato. Using a combination of histological and live-imaging techniques, together with spectroscopy, gene expression analysis and gene activation we provide important new insights into the pathogen-induced formation of vascular coatings. In particular, we show that ligno-suberin vascular coating and tyramine-derived HCAAs contribute to restriction of *R. solanacearum* in resistant tomato. In addition, we demonstrate that genes in the ligno-suberin-associated pathways can be explored to engineer resistance against *R. solanacearum* into commercial susceptible varieties of tomato.

## Results

### Resistant H7996 tomato restricts *R. solanacearum* colonization and induces a vascular coating response involving wall-bound phenolics

In order to understand the mechanisms underscoring restriction of *R. solanacearum* spread in resistant tomato varieties we used the resistant variety Hawaii 7996 (H7996) and compared it to the susceptible cultivar Marmande. In our assay conditions, most Marmande plants were wilted 10 days after inoculation with *R. solanacearum* GMI1000, while H7996 plants remained largely asymptomatic (Fig. 1A, S1A and (Planas-Marquès *et al*., 2019). Accordingly, bacterial loads in the taproot were drastically reduced in H7996 compared to Marmande, confirming the remarkable bacterial restriction ability of this cultivar (Fig. S1B and (Planas-Marquès *et al*., 2019)).

**Figure 1:**
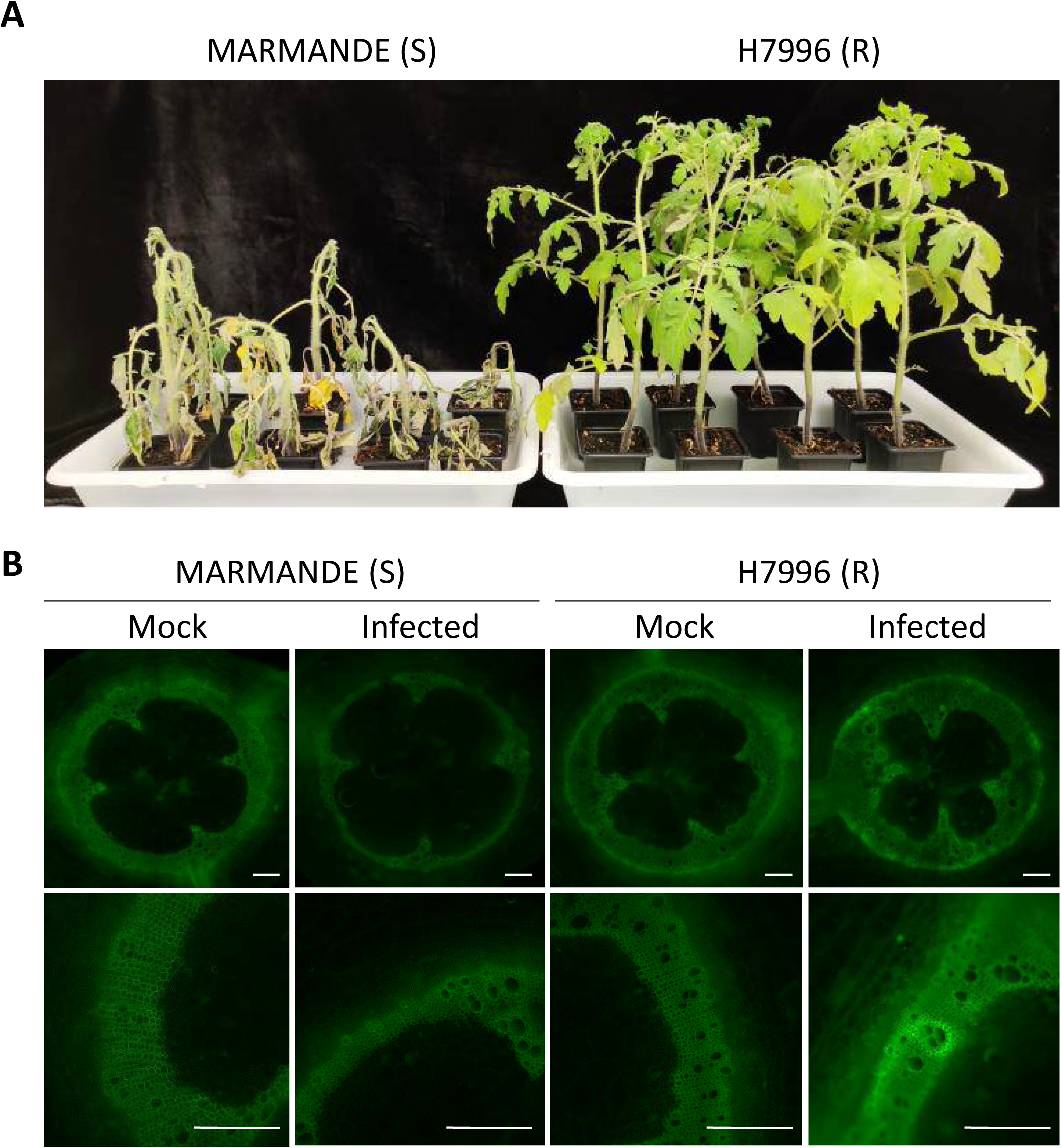
Resistant H7996 tomato restricts *R. solanacearum* colonization and induces a vascular coating response with wall bound phenolics. Susceptible (Marmande) and resistant (H7996), 5-week old tomato plants were inoculated through roots by soil-soak with ∼1×10^7^ CFU/ml of *R. solanacearum* GMI1000 and incubated at 28°C. **(A)** At 12 days post-inoculation (dpi) most Marmande plants showed severe wilting symptoms, whereas H7996 remained mostly symptomless. **(B)** Taproot cross-sections were obtained at 9 days post-infection (dpi). UV microscopy showed a strong autoflorescence signal emitted from the walls of vessels and surrounding parenchyma cells in infected H7996 plants compared to Marmande or the mock controls. Fluorescence signal in white was green colored. Images from a representative experiment out of 3 with *n*=5 plants per cultivar. Scale bar = 500 µm.

To identify defense-associated anatomical and/or physico-chemical modifications in H7996 after infection with *R. solanacearum* compared to Marmande we first analyzed ultraviolet (UV) autofluorescence of transverse taproot cross-sections, indicative of phenolic compounds (Donaldson, 2020). To focus on cell wall-deposited phenolic compounds, soluble phenolic compounds were removed with ethanol prior to observation as reported (Pouzoulet *et al*., 2013; Araujo *et al*., 2014). Infection with *R. solanacearum* induced a strong UV signal emitted from the walls of the vessels, and also from surrounding xylem parenchyma cells and tracheids in resistant H7996 (Fig. 1B). This enhanced autofluorescence was not observed in the susceptible variety Marmande nor in mock-treated samples (Fig. 1B).

### Spectroscopic analysis reveals *R. solanacearum*-induced deposition of suberin and accumulation of tyramine-derived amides in roots of resistant H7996 tomato and lignin structural modifications in roots of susceptible Marmande tomato

In order to decipher the composition of the cell wall-deposited compounds we used two complementary spectroscopic techniques: Fourier transform infrared spectroscopy (FT-IR) and two-dimensional heteronuclear single quantum correlation nuclear magnetic resonance (2D-HSQ NMR). Whereas FT-IR allows rapid analysis of the metabolic composition of a tissue at a given time (Türker-Kaya and Huck, 2017), 2D-HSQC NMR is considered one of the most powerful tools for plant cell wall structural analysis providing information on the composition and linkages in lignin/suberin polymers (Ralph and Landucci, 2010; Correia *et al*., 2020).

FT-IR confirmed that the most characteristic spectral features visible on the spectra of taproot vascular and paravascular tissue were connected with the presence of functional chemical groups of phenolic compounds (Fig. S2A and C). Calculation of relative absorbance ratios of the most diagnostic peaks showed a specific increase in phenolic compounds in resistant H7996 after infection with *R. solanacerum*, as can be seen by the higher phenolic –OH stretching value (≈ 3300 cm^-1^) when comparing to its mock control or susceptible Marmande plants (Fig. S2B).

To deepen our understanding of the compounds involved, 2D-HSQC spectra of infected or mock-treated taproots of H7996 and Marmande tomato plants were obtained and the main lignin and suberin substructures identified are shown in Fig. 2, while the chemical shifts of the assigned cross-signals are detailed in Table S1. Importantly, the aliphatic region of the 2D-HSQC spectra revealed that H7996 infected plants were more enriched in poly-aliphatic structures characteristic of suberin (magenta-colored signals), compared to its mock control (Fig 2A). Related to this, an olefinic cross-signal of unsaturated fatty acid structures (UF, δ_C_/δ_H_ 129.4/5.31), typical of suberin, was also found to be increased in the HSQC spectrum of the infected H7996 tomato. A rough estimate based on the integration of lignin and suberin HSQC signals, revealed that the suberin/lignin ratio in *R. solanacearum*-infected H7996 plants was doubled compared to mock-treated plants, evidencing an increase in suberin deposition as a consequence of the bacterial infection. Interestingly, signals compatible with feruloylamides (FAm_7_; δ_C_/δ_H_ 138.6/7.31) and with tyramine-derived amides (Ty in orange; δ_C_/δ_H_ 129.3/6.92, 114.8/6.64, 40.5/3.29 and 34.2/2.62) were exclusively found in the spectrum of infected H7996 plants, suggesting the presence of feruloyltyramine exclusively in these samples (Fig. 2A). Since tyramines have been found as structural components co-ocurring with suberin (Bernards *et al*., 1995; Bernards and Lewis, 1998), which generates physically and chemically resistant barriers (He and Ding, 2020), our results substantiate the hypothesis of suberin as an important defense element against *R. solanacearum* infection in resistant tomato plants. On the contrary, the 2D-HSQC spectra of the lignin/suberin fractions isolated from the Marmande variety did not display notable variations between mock and infected plants in the signals corresponding to suberin, tyramine-related structures nor feruloylamides (Fig 2A).

**Figure 2:**
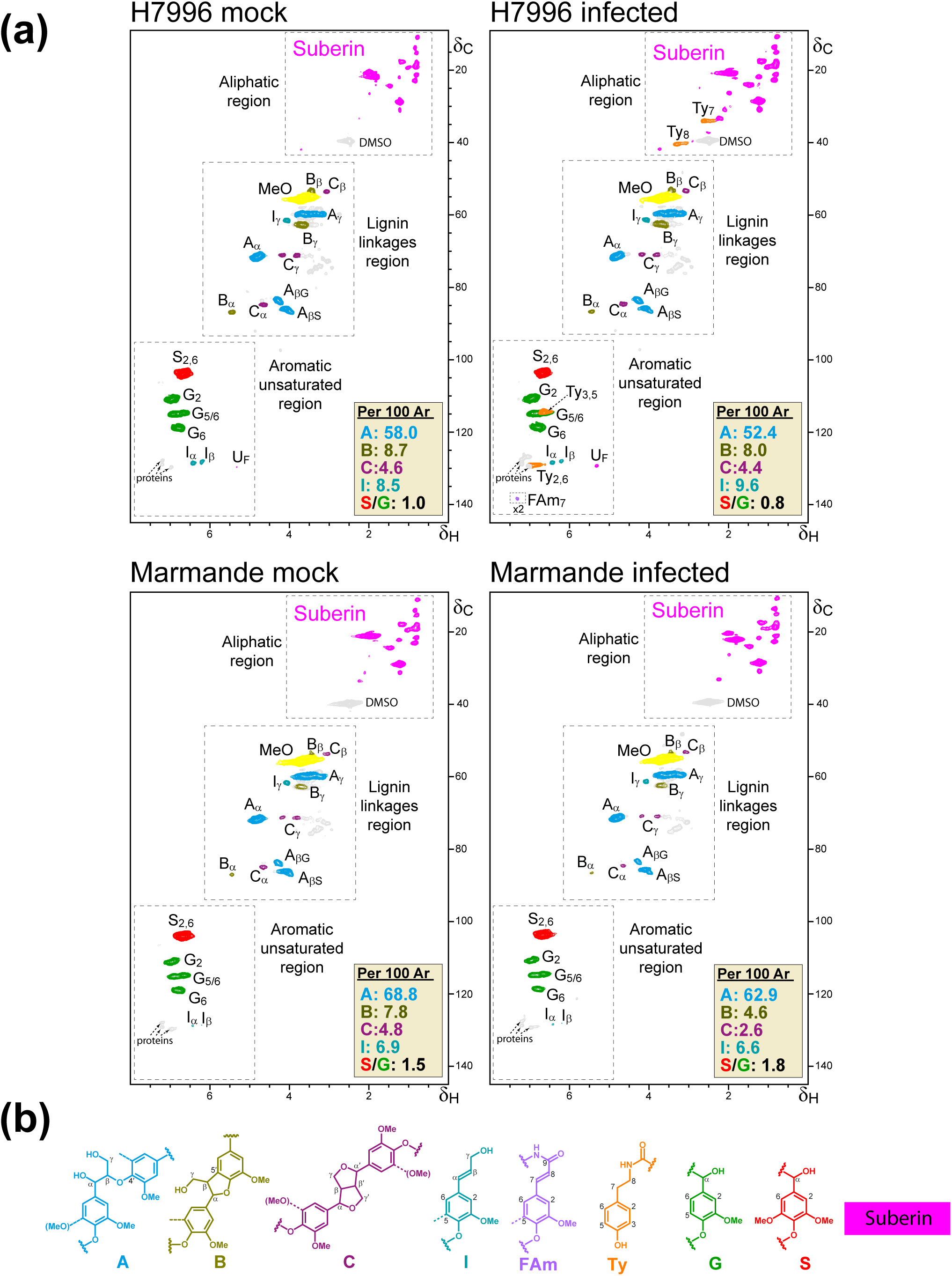
Feruloylamides, tyramine-derived amides and suberin-compatible compounds are specifically enriched in resistant H7996 tomato after infection with *R. solanacearum*. (A) 2D-HSQC NMR spectra of enzymatically isolated lignin/suberin fractions from mock-treated and *R. solanacearum*-infected taproots of H7996 and Marmande tomato plants. **(B)** Main lignin/suberin structures identified: β–*O*–4′ alkyl aryl ethers (A), β–5′ fenylcoumarans (B), β–β′ resinols (C), cinnamyl alcohols end-groups (I), feruloylamides (FAm), tyramine-derived amides (Ty), guaiacyl lignin units (G), syringyl lignin units (S), as well as unassigned aliphatic signals from suberin. The structures and contours of the HSQC signals are color coded to aid interpretation. ^1^H and ^13^C NMR chemical shifts of the assigned signals are detailed in Table S1. To detect FAm_7_ signal, the spectrum scaled-up to 2-fold (×2) intensity. The abundances of the main lignin linkages (A, B and C) and cinnamyl alcohol end-groups (I) are referred to as a percentage of the total lignin units (S + G = 100%).

Interestingly, 2D-HSQC NMR spectra also revealed significant structural modifications in the composition of lignin and the distribution of linkages in tomato plants after infection. Lignins with lower S/G ratios are more branched (condensed) and recalcitrant towards pathogen attack (Iiyama *et al*., 2020). Therefore, lignin in H7996, with an S/G ratio of 1.0 should be, a priori, more resistant than the lignin in Marmande plants (S/G ratio of 1.5). 2D-HSQC analysis revealed that the infection of susceptible Marmande plants resulted in an increase of the S/G ratio (from 1.5 to 1.8) and a clear reduction of all major lignin linkages (β–*O*–4′, β–5′ and β–β′; reduction in roughly 9%, 43% and 46%, respectively), evidencing that a lignin depolymerization process took place (Fig. 2A). In contrast, infected H7996 tomato roots displayed a slight decrease of the S/G ratio (from 1 to 0.8) (Fig. 2A), and only β–*O*–4′ linkages (the easiest to degrade in the lignin polymer) were significantly reduced (in roughly 10%), while the β–5′ and β–β′ were not so affected as in the case of Marmande plants. In this context, the major reduction in lignin linkages observed in Marmande after infection could explain, at least in part, its higher susceptibility to the pathogen.

### Histochemical analysis reveals the formation of structural vascular coatings containing suberin and ferulate/feruloylamide in resistant H7996 tomato roots in response to *R. solanacearum* infection

To confirm our spectroscopic data, we histochemically analyzed taproot samples of mock and infected H7996 and Marmande tomato plants. Observation of Phloroglucinol-HCl stained sections under brightfield microscopy (Wiesner staining) (Pomar *et al*., 2002; Pradhan Mitra and Loqué, 2014), showed that mock and infected H9776 (resistant) as well as mock Marmande (susceptible) samples showed a red-purple color characteristic of the reaction of phloroglucinol-HCl in vessels and fibers, indicative of lignin (Fig. 3A). In contrast, infected Marmande taproot sections exhibited reduced phloroglucinol-HCl staining, primarily concentrated in the xylem with less staining in the interfascicular fibers, suggesting a change in composition of xylem lignin upon infection, especially in the cell walls of fibers (Fig. 3A). This observation is in agreement with the structural changes specifically detected in the lignin structure of infected Marmande plants by 2D-HSQC NMR (Fig. 2A), which suggest lignin depolymerization and may partly underscore the high susceptibility of this tomato variety to *R. solanacearum*.

**Figure 3:**
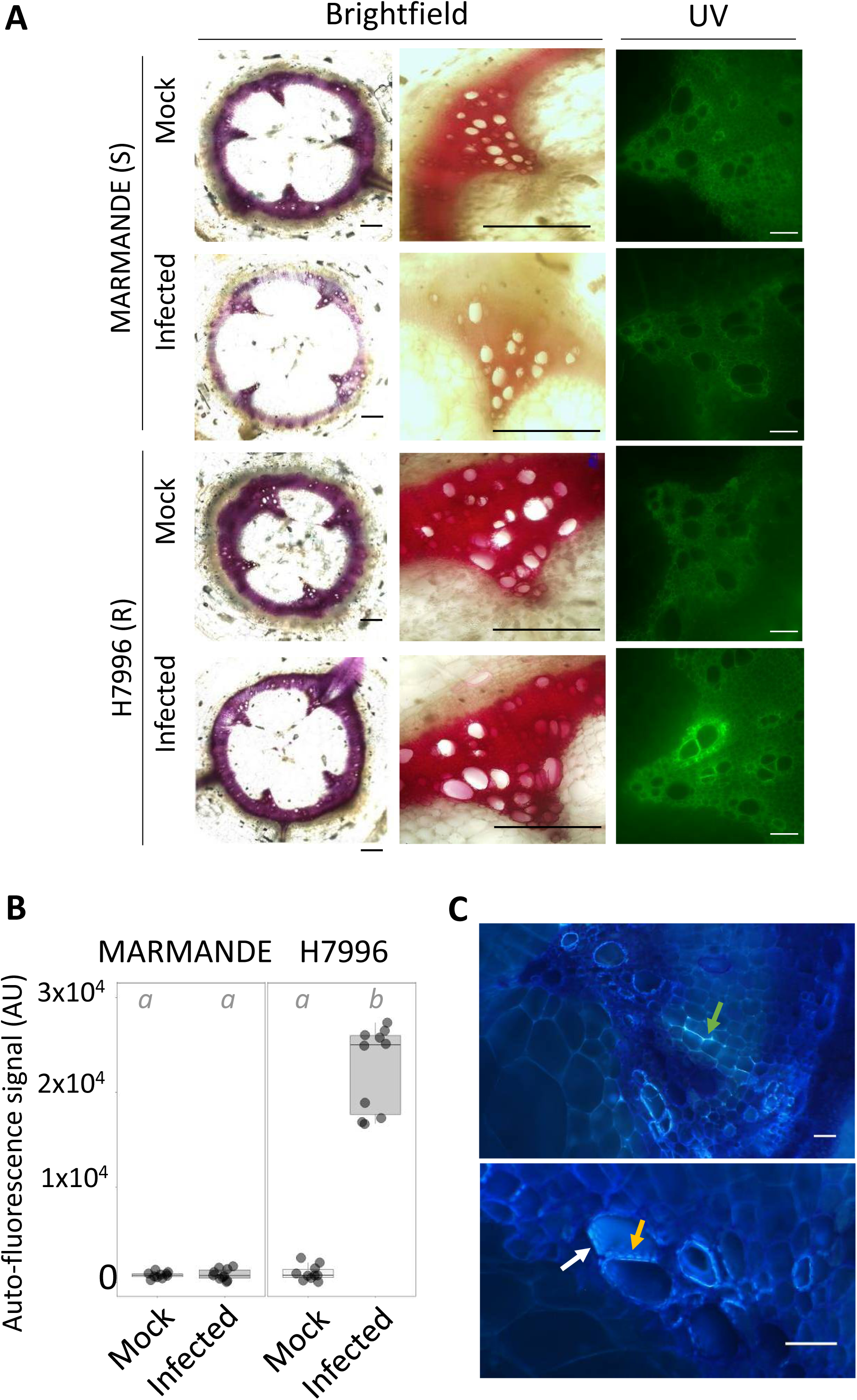
Resistant H7996 tomato shows vascular autofluorescence not-quenched with phloroglucinol and susceptible Marmande shows a decrease in phloroglucinol-HCl lignin signal. Susceptible (Marmande) and resistant (H7996) 5-week-old tomato plants were root-inoculated with a *R. solanacearum* GMI1000 strain at a concentration of ∼1×10^7^ CFU/ml or water mock. **(A)** Taproot cross-sections containing 10^5^ CFU g ^−1^ of *R. solanacearum* were stained with phloroglucinol-HCl and observed under UV to visualize other autofluorescent compounds different from lignin (not quenched with phloroglucionol- HCl) (left) and under brightfield to visualize lignin deposition (right). In infected H7996 strong UV autofluorescence could be observed in the walls of xylem vessels surrounding xylem parenchyma cells and tracheids, indicating reinforcement of walls of vascular tissue with phenolics formed *de novo* upon infection. In infected Marmande the red phlorogucinol stain was reduced especially in the intervessel areas. **(B)** The UV auto-fluorescence signal in (A) was measured using the LAS X Leica software after the Phloroglucinol-HCl treatment. **(C)** Detailed observation of infected H7996 xylem after the Phloroglucinol-HCl treatment shows the strong UV fluorescence concentrated in specific areas possibly corresponding to intervessel and vessel-parenchyma bordered pit membranes and/or pit chambers (yellow and white arrows, respectively). Fluorescence was also observed in parenchyma cells, specially enriched at intercellular cell corners (green arrow). (B) correspond to a representative experiment out of 3 each with n=6 plants per variety. Different letters indicate statistically significant differences (α=0.05, Fisher’s least significant difference test). (A) and (C) were representative images. Scale bars = 100 µm in (A, left), 500 µm in (A, right) and 50 µm in (C).

Ultraviolet (UV) illumination of phloroglucinol-HCl-stained samples allows quenching the autofluorescence from lignin and hence detect residual cell wall autofluorescence, which has been associated with suberin deposits (Baayen and Elgersma, 1985; Rioux *et al*., 1998; Pouzoulet *et al*., 2013). Under these conditions the increased autofluorescence observed in the vascular coating regions of infected H7996 tomato plants was not quenched in phlorogucionol-HCl stained samples (Fig. 3A, B). A more detailed observation revealed that this not-quenched autofluorescence was localized in specific regions compatible with (i) intervessel and vessel-parenchyma pit membranes or pit chamber walls and (ii) parenchyma coatings with fluorescent signals enriched in intracellular spaces (Fig 3C).

Since phenolic autofluorescence from walls that cannot be quenched by phloroglucinol-HCl treatment could be attributed to the suberin polymer (Biggs, 1984; Pouzoulet *et al*., 2013), we analyzed whether the pathogen-induced coating of vessels observed in H7996 correlated also with an increase in ferulates, a major suberin component. We performed KOH treatment of plant tissues, which specifically shifts the UV fluorescence of ferulate/feruloylamide to green, allowing its detection (Carnachan and Harris, 2000; Harris and Trethewey, 2010; Donaldson and Williams, 2018). UV autofluorescence of vascular coatings in response to *R. solanacearum* infection in resistant H7996 shifted from blue to a strong green color upon treatment with alkali (1N KOH) (Fig. S3A). This indicated that the *R. solanacearum*-induced xylem vasculature feruloylation was specific to resistant H7996, as the fainter blue autofluorescence observed in mock-treated resistant H7996 or susceptible Marmande tissues did not change to green at high pH in either early (Fig. S3A, B) or late (Fig. S3C) stages of infection.

To corroborate that the ferulate/feruloylamide accumulation in infected H7996 tomato was related with vascular suberization, we combined the ferulate-specific UV-alkali treatment described above with Sudan IV staining, which binds aliphatic components of suberin to produce a reddish-brown coloration. This revealed suberization in the taproot of *R. solanacearum*-infected H7996 plants, xylem vessel walls as well as the layers of vessels, parenchyma cells and tracheids in the immediate vicinity (reddish-brown signal from Sudan IV, Fig. 4). In the periphery of suberized cells, a green signal from UV-alkali was observed (Fig. 4), which may indicate ferulate/feruloylamide deposition indicative of a preceding stage towards suberization in this cell layer. In comparison, no positive Sudan IV or UV- alkali staining was detected in infected Marmande or mock-treated tomato plants. Together, suberized and feruloylated layers of parenchyma cells, vessels and tracheids might form a “suberization zone” creating a strong physico-chemical barrier to limit *R. solanacearum* spread from the colonized xylem vessel lumen.

**Figure 4:**
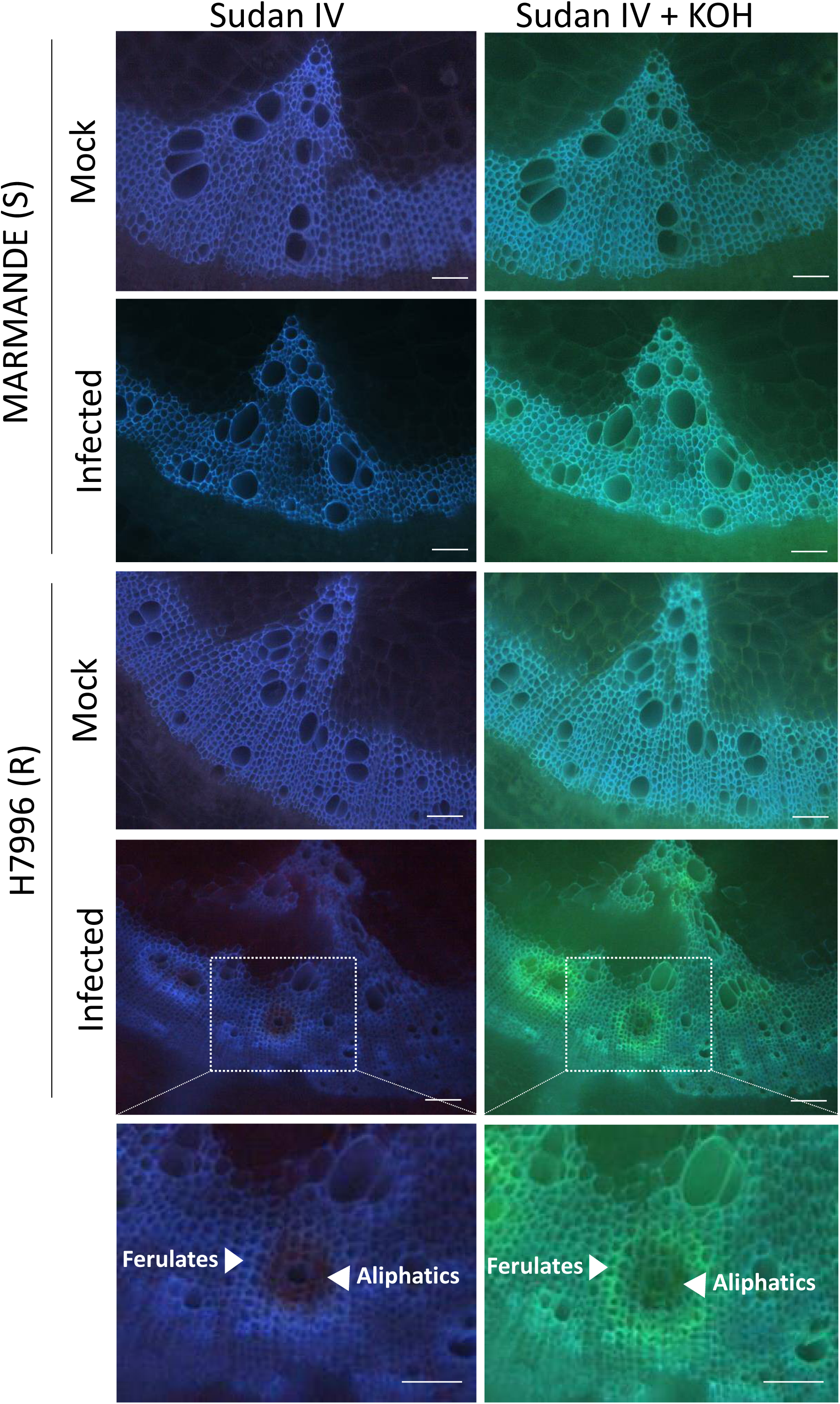
Resistant H7996 tomato shows cell wall ferulic acid and suberin deposition in restricted zones of vascular tissue upon *R. solanacearum* infection. Susceptible Marmande or resistant H7996 tomato plants were soil-inoculated with a ∼1×10^7^ CFU/ml suspension of *Ralstonia solanacearum* GMI1000 or mock-inoculated with water and incubated at 28°C. Cross-sections were obtained from taproot tissue containing 10^5^ CFU g^−1^ of *R. solanacearum*. Sections were stained with Sudan IV to visualize suberin aliphatics and subsequently treated with 1N KOH (pH above 10) to visualize ferulic acid bound to cell wall. Sudan IV positive staining (reddish-brown coloration) was observed around xylem vessels specifically in infected H7996, indicating accumulation of suberin aliphatics. Accumulation of ferulic acid bound to cell wall (blue-green coloration) appears also specifically in infected H7996 resistant tomato, surrounding sudan IV-stained areas. White arrowheads indicate the sites of accumulation of ferulates and aliphatic compounds. Representative images from one experiment out of three with *n*=6 plants each were taken.

### *R. solanacearum* infection activates the biosynthesis of aliphatic suberin precursors and feruloylamide, and aliphatic esterification of ferulic acid in the vasculature of resistant H7996

Since a differential accumulation of suberin-compatible compounds was specifically observed in infected H7996, we surmised that genes related to suberin and feruloylamide synthesis, as well as ferulic acid esterification to aliphatics may be upregulated in resistant tomato in response to *R. solanacearum* invasion. To test this hypothesis, we analyzed: i) expression of genes in the phenylpropanoid and suberin biosynthesis pathways, which provide the necessary precursors for the ligno-suberin heteropolymer; ii) the feruloyl transferase FHT (ASFT/HHT in Arabidopsis), which is involved in the formation of ferulate esters of fatty acyl compounds necessary to form suberin and soluble waxes (Molina *et al*., 2009; Gou *et al*., 2009; Serra *et al*., 2010); and iii) N-hydroxycinnamoyl transferases (*THT*), which are involved in the synthesis of HCAAs such as feruloyltyramine, previously found both as part of the lignin-like polymer and in the soluble phenolic fraction of some suberized tissues (Negrel *et al*., 1993; Schmidt *et al*., 1999).

Quantitative RT-PCR from taproot xylem vascular tissue of *R. solanacearum*- or mock-treated H7996 and Marmande plants showed specific upregulation of all genes analyzed from the suberin biosynthetic pathway in H7996 infected plants compared to the mock controls (Fig. 5, S4). These included essential suberin biosynthesis genes such as *CYP86A1* and *CYP86B1* (fatty acid oxidation), *FAR* (primary alcohol generation), *KCSs* (fatty acid elongases) and *GPAT5* (acylglycerol formation). In addition, feruloyl transferase FHT (ASFT/HHT in Arabidopsis), was also strongly upregulated in infected H7996 plants (Fig. 5 and S5). Regarding THT, in tomato we identified five putative homologs (Fig S6A), all induced by infection in the vascular tissue of H7996 (Fig. 5 and S6B). Among them, *SlTHT1-3* showed the strongest upregulation in H7996 after infection, although a slight upregulation could also be observed in Marmande (Fig. 5 and S6B). In comparison, *R. solanacearum* infection had only a modest effect in genes related to phenylpropanoid pathway as only upregulation was detected in the first enzyme of the pathway (PAL) (Fig. 5 and S7).

**Figure 5:**
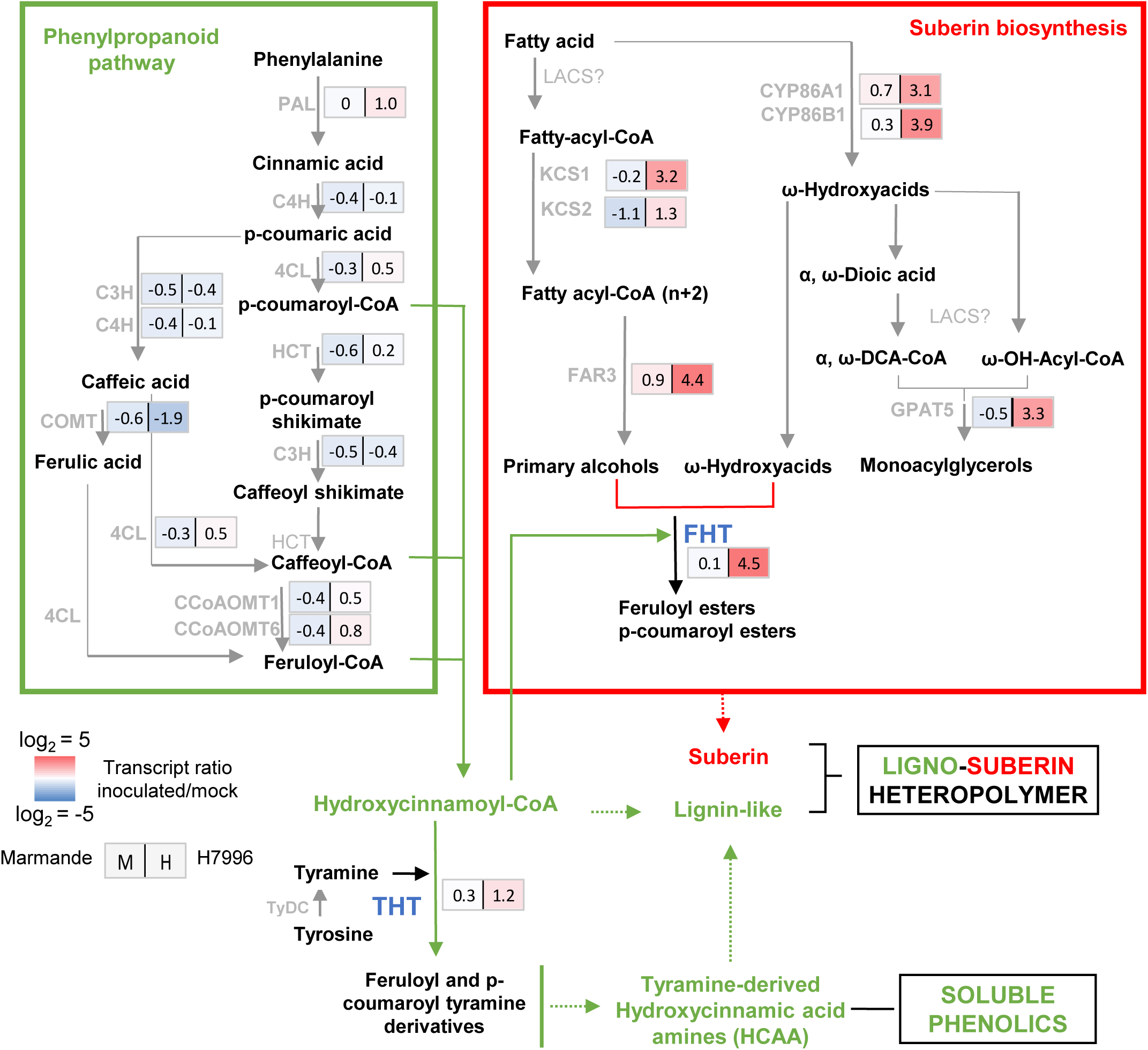
Genes of the ligno-suberin heteropolymer biosynthesis pathway are specifically induced in the xylem vasculature of resistant H7996 tomato upon *R. solanacearum*. The levels of expression of genes belonging to metabolic pathways relevant for suberin, lignin and feruloyltyramine and related amides biosynthesis were analyzed by qPCR of taproot vascular tissue in infected or mock-treated H7996 or Marmande tomato plants. Plants containing an *R. solanacearum* inoculum of 10^5^ CFU g^−1^ were selected and taproot xylem vascular tissue, comprising of metaxylems and surrounding parenchyma cells was collected for RNA extraction and cDNA synthesis. In parallel, xylem tissue was collected from mock plants. Heatmaps show log_2_ fold change RTA (relative transcript abundance) values of infected vs. mock for Marmande (left) and Hawaii (right). The tomato gene encoding for the alpha-subunit of the translation elongation factor 1 (*SleEF1 α*) was used as endogenous reference. Three biological replicates (n=3) were used, and taproots of 6 plants were used in each replicate. The scheme represents the phenylpropanoid and suberin biosynthesis pathways providing lignin-like and suberin precursors for the ligno-suberin heteropolymer. Abbreviations: PAL: Phenylalanine ammonia–lyase; C4H: Cinnamate–4–hydroxylase; C3H: Coumarate 3-hydroxylase; 4CL: 4–Coumarate–CoA ligase; HCT: Hydroxycinnamoyl–CoA shikimate/quinate hydroxycinnamoyl transferase; COMT: Caffeic acid 3-O-methyltransferase; CCoAOMT: Caffeoyl CoA 3-O-methyltransferase; CYP86A1 and CYP86B1: cytochrome P450 fatty acid ω-hydroxylases; KCS1/2: 3-ketoacyl-CoA synthase; FAR 1/3/4: Fatty acyl-CoA reductase; GPAT5: glycerol-3-phosphate acyltransferase 5; THT: Tyramine hydroxycinnamoyl transferase; TyDC: Tyrosine decarboxylase; FHT: feruloyl transferase. The question mark (?) denotes a hypothetical reaction.

Together, these data indicate that upregulation of genes involved in the formation of aliphatic suberin precursors, ferulic acid esterification to aliphatics (FHT) and production of HCAAs, such as feruloyltyramine (THT), constitute a very specific response of H7996 plants that takes place in the vasculature upon *R. solanacearum* infection. Further, these data are in agreement with NMR data of infected H7996, which showed a specific increase in insoluble fatty acid structures typical of suberin as well as the appearance of signals from structural tyramine-derived amides and feruloylamides (Fig. 2a). The higher expression of all these genes in H7996 upon infection may also result in the overproduction of soluble waxes and soluble HCAAs.

### Overexpression of *SlTHT1-3* in a susceptible tomato cultivar confers resistance to R*. solanacearum*

Based on our results, we set to determine whether overexpressing genes involved in ferulic acid esterification to suberin aliphatics and feruloylamide biosynthesis, such as *SlFHT* and *SlTHT1-3*, respectively, would increase resistance against *R. solanacearum* in a susceptible tomato background. First, we obtained transgenic tomato lines stably overexpressing *SlFHT* on a susceptible Marmande background (Fig. S8) and analyzed symptom progression and bacterial colonization. *SlFHT* overexpression lines showed a slight delay in disease progression (Fig. 6a) and moderately milder symptoms. The taproot and hypocotyl of *SlFHT* overexpressors displayed a slight reduction in bacterial loads after soil-soak inoculation in comparison to Wt tomato (Fig. 6b).

**Figure 6:**
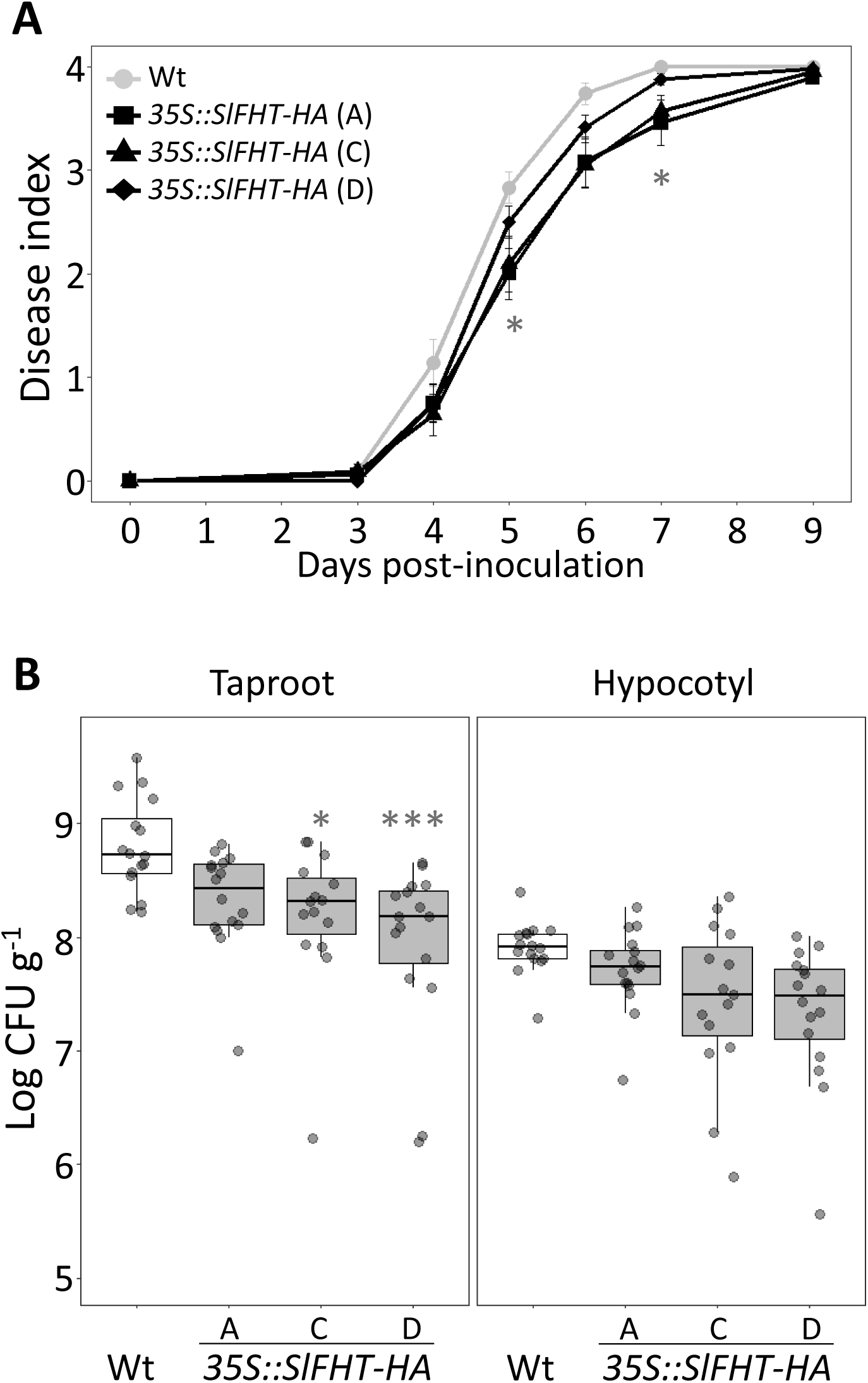
Overexpression of *SlFHT-HA* in susceptible tomato slightly restricts colonization by *R. solanacearum*. **(A, B)** A pathogenicity assay was performed comparing Wt and 3 independent *35S::SlFHT-HA* Marmande tomato lines (A, C and D) after infection with *R. solanacearum* GMI1000 lux reporter strain. Five-week-old plants were soil-soak inoculated with ∼1×10^7^ CFU/ml or mock and grown at 28°C. **(A)** Wilting progress was monitored by rating plants daily on a 0 to 4 disease index scale where 0 = healthy and 4 =100% wilted. Plotted values correspond to means ± standard error of 24 independent plants (n=24) from a representative experiment out of a total of 3. Asterisks indicate statistically significant differences between Wt and each of the *35S::FHT-HA* analyzed using a paired Student’s t-test (* p<0.05). **(B)** The level of *R. solanacearum* colonization in the taproot and hypocotyl was calculated as colony forming units per gram of fresh taproot tissue (CFUꞏg ^−1^) at 12 dpi. Data presented are of a representative experiment out of a total of 3 experiments. Asterisks indicate statistically significant differences between wild type and *35S::FHT-HA* tomato lines in a paired Student′s t-test (* corresponds to a p-value of p <0.05 and *** to p < 0.001).

Regarding *SlTHT1-3*, the corresponding tomato overexpressing line was readily available on a Moneymaker background (Campos *et al*., 2014). This line overaccumulates soluble HCAA such as feruloyltyramine. Using this *SlTHT1-3* overexpressing line and the corresponding Moneymaker wild type, we performed a variety of *R. solanacearum* infection assays. As expected, the Moneymaker tomato cultivar showed similar susceptibility to *R. solanacearum* as Marmande (Fig. 1a, b and Fig. 7a, b). In contrast, overexpression of *SlTHT1-3* resulted in a dramatic increase of resistance against *R. solanacearum,* with disease progressing remarkably slower in this line compared to the Wt Moneymaker (Fig. 7a, b). Importantly, bacterial loads were significantly lower in the taproot and hypocotyl of the *SlTHT1-3* overexpressor after soil inoculation in comparison to Wt tomato (Fig. 7c). Similarly, direct leaf inoculation also showed severe bacterial growth restriction in the *THT1-3* overexpressing line (Fig. S9a). Further, we monitored the colonization patterns of a *R. solanacearum* GFP reporter strain after stem inoculation of the *SlTHT1-3* overexpressing line compared to Wt. In transverse stem cross-sections of 6 dpi plants, bacteria stayed confined near the inoculation point in the *35S::SlTHT1-3* line whereas they spread unrestrictedly in susceptible wild type stems from the inoculation point and at least 3 cm up and downwards (Fig. 7d and S9b). Quantification of the GFP signal confirmed that bacterial growth was drastically reduced in *SlTHT1-3* overexpressing plants in comparison to Wt (Fig. 7e). Together, our data clearly show that *StTHT1-3* ectopic expression provides a very effective resistance mechanism against *R. solanacearum* - potentially mediated by accumulation of elevated amounts of HCAAs such as feruloyltyramine-, which drastically restricts vascular colonization, preventing bacterial spread and blocking the onset of disease.

**Figure 7:**
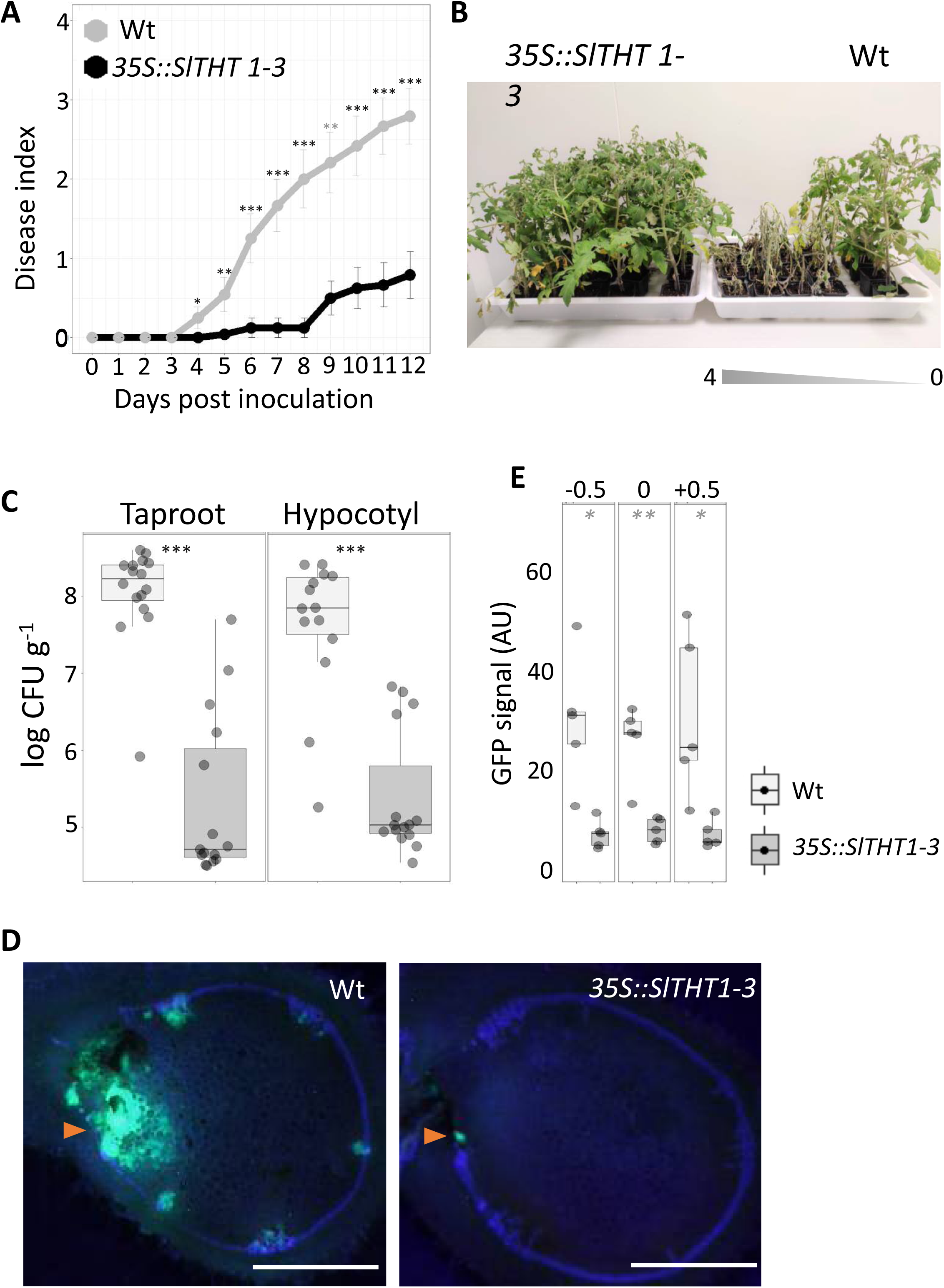
Overexpression of *SlTHT1-3* in susceptible tomato confers resistance to *R. solanacearum*. **(A, B)** A pathogenicity assay was performed comparing Wt and *35S::SlTHT1-3* tomato lines (Moneymaker background) after infection with *R. solanacearum* lux reporter strain of GMI1000. Five-week-old plants were soil-soak inoculated with ∼1×10^7^ CFU/ml and grown at 28°C. **(A)** Wilting progress was monitored by rating plants daily on a 0 to 4 disease index scale where 0 = healthy and 4 =100% wilted. Plotted values correspond to means ± standard error of 24 independent plants (n=24) from a representative experiment out of a total of 3. Asterisks indicate statistically significant differences between Wt and *35S::SlTHT1-3* using a paired Student’s t-test (* p<0.05, ** p<0.01 and *** p<0.001). **(B)** Pictures were taken 12 days post-infection. Wt plants were arranged according to the degree of symptom severity (from 4 to 0). **(C)** Transgenic *35S::SlTHT1-3* tomato significantly restricted *R. solanacearum* colonization in both the taproot and hypocotyl to Wt. Five-week-old tomato plants were root-inoculated with a *R. solanacearum* GMI1000 luciferase reporter strain at a concentration of ∼1×10^7^ CFU/ml or water mock. The level of *in planta* colonization by *R. solanacearum* was calculated as colony forming units per gram of fresh taproot tissue (CFUꞏg ^−1^) at 12dpi. Box-and-whisker plots show data from a single representative experiment out of 3 (n =14 to 16). **(D)** Transverse stem cross-sections of Wt and transgenic *35S::SlTHT1-3* tomato lines were imaged under a confocal microscope 6 days after infection with a *R. solanacearum* GMI1000 GFP reporter strain. *R. solanacearum* at a concentration of 10^5^ CFU ml ^−1^ was injected directly into the xylem vasculature of the first internode thorough the petiole. Representative images of *R. solanacearum* colonization progress at the point of inoculation are shown. **(E)** Mean green fluorescence of the GFP signal emitted from *R. solanacearum* at cross-sections obtained as described in (D) at the point of inoculation (0), below the point of inoculation (-0.5 cm) and above the point of inoculation (+0.5 cm) was measured using ImageJ. Data from a representative experiment out of a total of 3, with *n*=5 plants per condition. Asterisks indicate statistically significant differences between wild type and *35S::THT1-3* tomato plants in a paired Student′s t-test (* corresponds to a p-value of p <0.05, ** to p < 0.01 and *** to p < 0.001).

## Discussion

### Ligno-suberin deposits in vascular cell walls and feruloyltyramine accumulation acts as a resistance mechanism restricting *R. solanacearum* colonization in resistant tomato

The root xylem vasculature is one of the first sites of multiplication of *R. solanacearum* inside the host (Vasse *et al*., 1995; Álvarez *et al*., 2010; Digonnet *et al*., 2012). Colonization of the xylem vasculature is critical, as in this particular tissue the pathogen multiplies and moves vertically to the stem alongside the xylem fluid. In susceptible hosts, the pathogen also spreads horizontally from colonized vessels to the healthy neighboring tissues, including vessels and surrounding parenchyma cells (Nakaho *et al*., 2000). To facilitate this process *R. solanacearum* secretes an array of cell wall degrading enzymes (Liu *et al*., 2005; Lowe-Power *et al*., 2018) In parallel, plant cell walls also act as dynamic barriers against pathogens, acting as first line of defense by undergoing remodeling or strengthening upon pathogen recognition (Underwood, 2012). However, the precise role of cell walls in defense responses is far from being understood and has been mostly studied in the leaves.

In our study, resistant tomato (H7996) was observed to react aggressively to *R. solanacearum* infection by reinforcing the walls of vessels and the surrounding parenchyma cells with phenolic deposits, which can be observed as UV autofluorescence (Fig. 1). An increase in autofluorescence had been previously reported in another resistant tomato variety, LS-89, although its composition was not precisely defined (Ishihara *et al*., 2012). Histochemical analysis of vascular coatings in resistant tomato upon *R. solanacearum* infection showed that the strong UV autofluorescence emitted from xylem vessel walls and the surrounding parenchyma cells observed in resistant H7996 against *R. solanacearum* (Fig. 1) could not be quenched by phloroglucinol-HCl, suggesting the presence of suberin deposits (Fig. 3) (Biggs, 1984; Rittinger *et al*., 1986; Pouzoulet *et al*., 2013). Detailed observation revealed that this suberin-associated autofluorescence was prominent in vessel-parenchyma and intervessel pit membranes and/or chambers and in parenchyma intercellular spaces (Fig. 3C). In line with this, previous reports using TEM showed thickening of the pit membranes accumulating electron dense material in tomato plants resistant to *R. solanacearum* (Nahako *et al*., 2000 and 2004). The suberin nature of these coatings was further supported by the positive Sudan IV staining of vessels and surrounding parenchyma cells of H7996 roots upon infection (Fig. 4). These results are in agreement with the previous suberin coatings detected in tomato plants resistant to *Verticillium albo-atrum*. Upon infection or after treatment with the suberin hormone inducer abscisic acid, suberin was chemically detected in petioles (Robb *et al*., 1991), and suberin as well as lignin coatings, were both deposited in intercellular spaces between parenchyma cells adjoining a xylem vessel or infusing and occluding pit membranes coatings (Robb *et al*., 1991; Street *et al*., 1996). Besides, inhibition of the phenylpropanoid pathway by blocking PAL enzyme inhibited the formation of both lignin and suberin coatings (Street *et al*., 1996), in agreement with the ferulic acid requirement to correctly deposit suberin (Andersen *et al*., 2021) and reinforcing our observations of the presence of a ferulate/feruloylamide-derived polymer detected in H7996 *R. solanacearum* KOH-treated samples observed under UV light (Fig. 4). In line with this, 2D-HSQC NMR data of resistant H7996 tomato vascular tissue revealed the presence of tyramine-derived amides and feruloylamides incorporated into the cell wall and also an enrichment in poly-aliphatic structures characteristic of suberin (Fig. 2). An increase in feruloyltyramine has been detected associated with suberization (Graça, 2015; Legay *et al*., 2016; Figueiredo *et al*., 2020) and is compatible with the lignin coatings seen in conjunction with suberin coatings in *V. albo-atrum* infected tomatoes (Robb *et al*., 1991).

Interestingly, in the periphery of the suberized (Sudan IV-stained) layers surrounding the vasculature after infection in resistant tomato we could observe cells with intense accumulation of phenolics where Sudan IV did not bind. In these peripheral areas, the strong blue-to-green color conversion upon alkali treatment (Carnachan and Harris, 2000; Harris and Trethewey, 2010) revealed the presence of wall-bound ferulic acid, potentially derived from ester hydrolysis of suberin aliphatics (ferulates). However, since amides can also be hydrolyzed as esters in basic solution (Robert and Caserio, 1977), ferulic acid could alternatively be derived from wall-bound feruloylamides, such as the feruloyltyramines shown to accumulate in resistant tomato by 2D-HSQC NMR (Figure 2A) (Fig. 4, S3), which may act as further reinforcements against the pathogen. Importantly, in tomato vasculature infected with *V. albo-atrum* lignin deposits were detected preceding those of suberin (Robb *et al*., 1991).

Beyond histochemistry and spectroscopic signature detections of suberin, lignin and ether linked feruloyltyramine or related amides, further evidence supporting the nature of these ligno-suberin coatings as responsible of the resistance observed in H7796 to *R. solanacearum* was unequivocally provided transcriptionally using transcriptional gene markers. Tissues undergoing suberization have to go through a complex genetic and metabolic reprogramming involving a network of metabolic pathways, in order to produce the precursors of the polymer and subsequently their polymerization into the matrix (Lashbrooke *et al*., 2016). Transcriptional reprogramming associated to suberin biosynthesis was clearly observed in the root vascular tissue of resistant H7996 tomato upon infection with *R. solanacearum*. Specific upregulation in the xylem of resistant tomato of genes that are specific and key for suberin monomer biosynthesis was observed, including KCS elongases, FAR reductases, CYP86 ɷ-hydroxylases, GPAT5 acyltransferase and FHT/ASFT feruloyltransferase (Fig. 5). In contrast, only moderate differences were found in transcripts of phenylpropanoid pathway genes. Interestingly, PAL, which showed modest upregulation in resistant H7996, had been previously defined as a rate-limiting enzyme of phenylpropanoid pathway (Faragher and Brohier, 1984; Howles *et al*., 1996). Considering this, the observed upregulation could provide more tyramine and feruloyl-CoA, which together with the upregulation of *THT* would be in agreement with the increased presence of feruloyltyramine detected by 2D-HSQC NMR (Figure 2A). The slight upregulation in resistant H7996 and downregulation in susceptible Marmande of other phenylpropanoid pathway genes upon infection corroborates previous reports (Ishihara *et al*., 2012).

2D-HSQC NMR also revealed differences in the composition and structure of lignin between resistant and susceptible tomato cultivars after infection. The amounts and the level of lignin of a particular tissue affect wall strength, degradability and pathogen resistance (Mnich *et al*., 2020) and phenolic reinforcements in the xylem vasculature can act as a shield against pathogen-derived metabolites such as toxins and enzymes, and make water and nutrients inaccessible for pathogens, thereby impeding their growth (Araujo *et al*., 2014). Susceptible Marmande, which do not develop ligno-suberized coatings, presented predominance of S-type lignin, while the lower S/G ratio observed for the infected H7996 resistant cultivar, revealed a higher presence of a G-type lignin, which has also been related with the lignin accompanying suberized tissues (Graça, 2015). S-lignin is relatively unbranched and has a lower condensation degree than G lignin, which is more difficult to hydrolyze because it contains a higher proportion of condensed carbon-carbon linkages (Novaes *et al*., 2010). The observed lignin structural differences between varieties after infection were corroborated by phloroglucinol-HCl staining observed under bright field: in H7996 the G lignin together with the higher cross-linking resulted in strong red-purple positive staining, while S lignin, less cross-linking of cell wall polymers and a certain degree of degradation of the vascular root tissue resulted in less reaction to the stain (Kutscha and Gray, 1972; Hao *et al*., 2014) (Fig. 2 and 3). Together, these data indicate that i) under basal conditions the two tomato varieties used in this study display differences in the composition and structure of lignin and ii) *R. solanacearum* infection affects very differently the lignin fraction in the two varieties: resistant H7996 tomato shows only a slight decrease in the S/G ratio that may be linked to an accumulation of the ligno-suberin heteropolymer, while susceptible Marmande undergoes pronounced depolymerization that correlates with a decrease in phloroglucinol-HCl staining (Fig. 3A). Although the *R. solanacearum* has not been shown to be able to specifically depolymerize lignin, the pathogen secretes enzymes that can degrade cell wall polysaccharides and could participate in the observed Marmande stem collapse phenotype (Fig. 1A). In resistant H7996, on the other hand, vascular suberin-containing coatings would allow to create a hydrophobic barrier to prevent enzymes from accessing the cell wall substrates and at the same time create reinforcements, leading to resistance to the pathogen. The fact that these reinforcements are rich in tyramine or feruloyltyramine as shown by HSQC-NMR, may further reinforce the structural barrier formed in H7996 in response to R. solanacearum, providing rigidity, hampering cell wall digestibility by the pathogen’s hydrolytic enzymes (Macoy *et al*., 2015; Zeiss *et al*., 2020).

Overall, our data indicate that vascular coating with wall-bound ligno-suberized compounds may restrict horizontal spread of the bacterium at early stages of bacterial colonization (starting at ∼10^5^ CFU g ^−1^ taproot tissue), before the plant shows any visible wilting symptom (Fig 1). In comparison, susceptible tomato (Marmande) is either not able to induce such vascular coating upon *R. solanacearum* infection or induces a very weak and late response (Figs. 1, 3), potentially predisposing its vascular walls to disruption by the pathogen’s cell wall degrading enzymes. In absence of ligno-suberin reinforcements, the bacterium multiplies and colonizes abundantly moving out from vessel lumen into surrounding parenchyma cells and apoplastic spaces. Based on the above observations, we propose a model whereby resistant H7996 tomato undergoes vascular suberization upon *R. solanacearum* infection as follows (Fig. 8). When reaching the xylem vessels of resistant H7996, *R. solanacearum* multiplies and tries to invade the surrounding healthy vessels and parenchyma cells by degradation of the xylem pit membranes and walls. Resistant tomato plants respond to vascular invasion by the pathogen depositing feruloyltyramine and other HCAA-tyramine derived compounds, and suberin. These deposits would block the pit membrane access and serve as coatings of the vessel walls and parenchyma cells present in the immediate vicinity of colonized vessels, compartmentalizing the infection. These ligno-suberized layers of parenchyma cells, vessels and tracheids together form a “zone of ligno-suberization” creating a strong physico-chemical barrier to limit *R. solanacearum* spread from colonized xylem vessel lumen.

**Figure 8:**
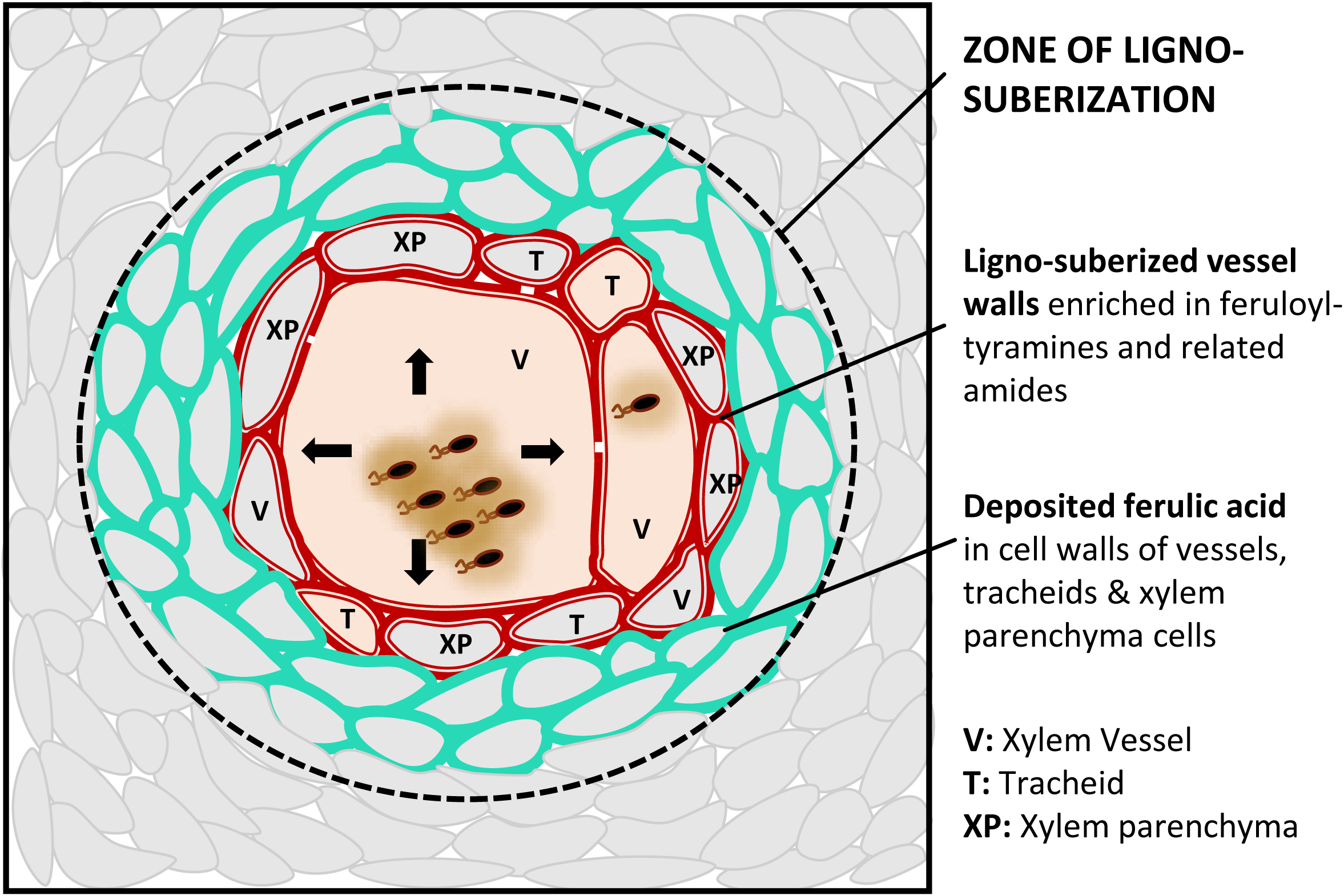
Schematic representation of the vascular ligno-suberization process potentially taking place in infected vessels of resistant H7996 tomato upon *R. solanacearum* infection. Colonization of the vasculature by *R. solanacearum* in resistant tomato plants induces a ligno-suberization process in the walls of the infected vessel (V) and of the adjacent tracheids (T) and parenchyma cells (XP) (red). The lignin-like polymer accompanying suberin would be enriched in structural feruloyltyramine and related amides. The signal of structural ferulic acid (ester or as amide) would extend to the walls of peripheral parenchyma cells, vessels and tracheids (green), indicating a stage preceding suberization or a final layered pattern, still to be resolved. Together, the red and green areas, would form a “zone of ligno-suberization” (black dashed line) potentially creating a physico-chemical barrier to limit *R. solanacearum* spread from the colonized xylem vessel lumen.

### Engineering tomato resistance against *R. solanacearum* by inducing the tyramine-HCAA pathway

Considering the observed accumulation of lignosuberin and cell wall-linked feruloyltyramine in resistant H7996 tomato in response to *R. solanacearum* infection, we sought to understand the implications of overexpressing genes involved in the synthesis of these compounds in susceptible tomato cultivars upon *R. solanacearum* infection. We focused on FHT and THT and because their corresponding transcripts are upregulated in the xylem vasculature of resistant tomato upon *R. solanacearum* infection (Fig. 5) and they are the enzymes related with the synthesis of suberin ferulates and ether linked feruloyltyramine, respectively.

*SlFHT* overexpression had a small effect on the responses of susceptible tomato against *R. solanacearum*, showing a slight delay in wilting symptoms together with a slight decrease of bacterial loads in the plant (Fig. 6). In resistant H7996 tomato, *SlFHT* was highly induced in and around the vasculature upon *R. solanacearum* expression, and this was accompanied by a strong upregulation in fatty acid biosynthesis genes (Fig. 5, S5), which provide key precursors to form suberin. The fact that increasing the levels of FHT in Marmande does only result in a marginal increase in resistance might be linked to a shortfall of aliphatic precursors in this variety (Fig. 5), which constrain a subsequent increase in suberin synthesis.

In contrast, transgenic tomato overexpressing *SlTHT1-3* in a susceptible Moneymaker background was highly resistant to *R. solanacearum*. Wilting symptoms and *in planta* bacterial loads were drastically reduced and colonization was dramatically restricted (Fig. 7). Importantly, this transgenic line was previously shown to accumulate elevated amounts of soluble HCAAs such as feruloyltyramine upon infection with the bacteria *Pseudomonas syringae* pv. *tomato* (*Pto*) (Campos *et al*., 2014). Disease susceptibility towards *Pto* was slightly reduced in the *SlTHT1-3* overexpressing lines (Campos *et al*., 2014). The fact that overexpressing *SlTHT1-3* in tomato confers strong resistance against *R. solanacearum*, but only a marginal increase in resistance against *Pto* indicates that enhanced production of tyramine-derived HCAAs constitute an important defense strategy against vascular pathogens, while for foliar pathogens other mechanisms are in place. There are several evidences that feruloyltyramine exhibit antimicrobial activity (Fattorusso *et al*., 1999; Novo *et al*., 2017) and that they can be involved in plant priming or an adaptive strategy where plants are in a physiological state with improved defensive capacity (Zeiss *et al*., 2021). These tyramine-derived HCAAs overproduced in *SlTHT1-3* overexpressing tomato lines may interfere with *R. solanacearum* colonization by i) becoming incorporated into the vascular and perivascular cell walls, providing a stronger cross-linking and restricting the movement of the pathogen inside the plant and/or ii) remaining soluble and acting as direct antimicrobial agents against the pathogen. The fact that *R. solanacearum* possesses a hydroxycinnamic acid degradation pathway, and mutants that cannot degrade hydroxycinnamic acids are less virulent on tomato (Zhang *et al*., 2019) clearly underscores the importance of HCAAs in the arms race taking place in this pathosystem.

In conclusion, we have provided evidence of the formation of a “ligno-suberization zone” enriched in ether linked feruloyltyramine and possibly related amides as an effective strategy to confine *R. solanacaerum* into infected vessels of resistant tomato plants, preventing horizontal spread of the pathogen into healthy tissues and delaying disease symptoms. Resistance against *R. solanacearum* can be attained in susceptible tomato background by stably overexpressing *THT*, potentially contributing to the formation of a lignin physico-chemical barrier and/or through a direct anti-microbial effect. Still, many questions remain to be answered. In the future, it will be interesting to investigate the contribution of HCAAs and suberin to resistance against the pathogen, the mechanisms whereby *R. solanacearum* perception leads to the formation of a ligno-suberin coatings around the vasculature in resistant tomato varieties. Increasing the spatio-temporal resolution of the tomato-*R. solanacearum* interaction will be instrumental to reach a deeper insight into structural resistance mechanisms. Also, since vascular confinement has been reported in different plant species as a means of resistance against various vascular wilt pathogens (De Ascensao and Dubery, 2000; Martín *et al*., 2008; Xu *et al*., 2011; Sabella *et al*., 2018), the level of conservation of vascular ligno-suberin deposition as a constituent of vascular coatings and part of a resistance mechanism remains to be determined.

## Materials and Methods

### Plant materials and growth conditions

The tomato (*Solanum lycopersicum*) varieties used in this study were the susceptible commercial variety Marmande and the quantitatively resistant public open-pollinated breeding line Hawaii 7996. We also used the tomato variety Moneymaker wild-type and *35s::THT 1-3,* generated by Campos *et al*., (2014). Seeds were germinated and plants were grown in pots consisting of soil (Substrate 2, Klasmann-Deilmann GmbH) mixed with perlite and vermiculite (30:1:1) in controlled growth chambers at 60% humidity and 12 h day/night with light intensity of 120–150 µmolꞏm^−2^ꞏs^−1^. Temperature was set at 27°C when using LED lighting and at 25°C when using fluorescent lighting.

### *Ralstonia solanacearum* strains and growth conditions

All assays in tomato were performed using *R. solanacearum* GMI1000 strain (Phylotype I, race 1 biovar 3). Luminescent and fluorescent reporter strains of *R. solanacearum* GMI1000 were used in the study containing constructs *PpsbA::LuxCDABE* and *PpsbA::GFPuv,* respectively (Cruz *et al*., 2014; Planas-Marquès *et al*., 2019).

### DNA constructs

For generation *35S::FHT-HA* construct the *FHT* (Solyc03g097500) coding sequence was amplified from tomato H7996 cDNA using the forward primer (part7FHTF1), having a flanking *Sma*I restriction enzyme digestion site at 5’ end and reverse primer (part7FHTHAR1), including the sequence of hemagglutinin (HA) epitope tag and a BamHI restriction enzyme digestion site at the 5’ end. The amplified product was cloned into the pJET1.2/blunt cloning vector using CloneJet PCR cloning kit (Thermofisher) and then digested by *Sma*I and *Bam*HI. The digested products were purified using NZYGelpure (Nzytech) followed by ligation into the pART7 and later to pART27 vector (Gleave, 1992).

### Stable transformation of tomato

*35S::FHT-HA* were transformed into Marmande. For this, the construct was transformed into *Agrobacterium tumefaciens* strain C58C1. *A. tumefaciens* was used for co-culture with tomato cotyledons. Explant preparation, selection, and regeneration followed the methods described by (Mazier *et al*., 2011). Transformants were selected on kanamycin-containing medium. Accumulation of FHT-HA protein was assayed by immunoblot with a monoclonal HA antibody (GenScript).

### Bacterial inoculation in plants

Four to five week-old tomato plants were inoculated through roots with *R. solanacearum* using the soil drenching method. For this, roots were wounded by making four holes in the corners of the pot with a 1 ml pipette tip and inoculated with a to 1×10^7^ CFU ml^-1^ (OD_600_ = 0.01) suspension of bacteria (Planas-Marquès *et al*., 2018). Inoculated plants were kept in a growth chamber at 27°C. For tomato leaf infiltration, plants were vacuum-infiltrated by submerging the whole aerial part in a ∼10^5^ CFU ml^-1^ (OD_600_ = 0.0001) *R. solanacearum* suspension as described in Planas-Marquès *et al*., (2018). For inoculation directly onto the stem vasculature, 10 µl (5 µl at a time) of 10^5^ CFU ml^-1^ (OD_600_ = 0.0001) *R. solanacearum* suspension was placed at the node of the petiole and pin-inoculated using a sterile 0.3×13 mm needle (30G×½″, BD Microlance, Becton Dickinson).

### *R. solanacearum* pathogenicity assays and quantification of bacterial growth *in planta*

Infected plants were scored for wilting symptoms using a scale from 0 to 4: 0=healthy plant with no wilt, 1=25%, 2=50%, 3=75%, and 4=100% of the canopy wilted as described by Planas-Marquès *et al*., (2019). The relative light units per second (RLUꞏs^−1^) readings were converted to CFUꞏg^−1^ tissue as described in Planas-Marquès *et al*., (2019). For bacterial colonization assays using GFP reporter strain, transverse stem cross-sections were made at the inoculation point as well as at a distance of 0.5 cm, 1 cm, 2 cm and 3 cm in both upward and downward direction, using a sterile razor blade. The sections were photographed using an Olympus SZX16 stereomicroscope with a UV fluorescent lamp (BP330-385 BA420 filter) and equipped with a DP71 camera system (Olympus). Quantification of mean green fluorescence from xylem vascular ring and pith parenchyma was done using ImageJ software (Planas-Marquès *et al*., 2019). For leaf *in planta* multiplication assays, 3 leaf discs of 0.8 cm^2^ size were homogenized in 200 µl of sterile distilled water. CFU cm^-2^ leaf tissue were calculated after dilution plating of samples with appropriate selection antibiotics and CFU counting 24 hours later.

### Histological methods

Taproots of *R. solanacearum-*soil drench inoculated or water-treated plants were used for obtaining thin transverse cross-sections with a sterile razor blade. Inoculated plants were either sectioned at 9 dpi or when bacterial colonization level reached 10^5^ CFU g^-1^ taproot tissue, as indicated in the figure legend where only H7996 sections showed a localized browning at one xylem pole indicative of infection and defense reactions at xylem. Sections were kept in 70 % ethanol at room temperature for 5-7 days and examined using fluorescence microscopy using a Leica DM6B-Z microscope under UV illumination (340-380 nm excitation and 410-450 nm barrier filters). Autofluorescence emitted from phenolic deposits was recorded using a Leica-DFC9000GT-VSC07341 camera and the signal was pseudo-colored green.

Sections were also stained with a Phloroglucinol-HCl solution (100 mg phloroglucinol in 8 ml of ethanol 95% and 8 ml of hydrochloric acid 37 %) for the detection of lignin and observed under bright field (Pomar *et al*., 2004). Photographs were taken with a DP71 Olympus color digital camera. Since Phloroglucinol binds to lignin and quenches the autofluorescence emitted from it (Martín *et al*., 2005), the cross-sections were then observed under UV microscopy to detect the remaining autofluorescence which would correspond to non-lignin phenolic sources such as suberin (Pouzoulet *et al*., 2013). A Leica-DM6B-Z microscope (340-380 nm excitation and 410-450 nm barrier filters) was used. Autofluorescence was recorded using a Leica-DFC9000GT-VSC07341 digital camera and the signal was pseudo-colored green. Detailed observations (Fig. 3c) were done using a Olympus AH2 Vanox-T microscope, exciting at 330-380 nm and collecting emission wavelengths from 420 nm. Images were recorded using the Olympus XC-50 color digital camera.

Auto-fluorescence from ferulic acid bound to the cell wall shows a pH-dependent blue to green color conversion (Harris and Trethewey, 2010; Carnachan and Harris, 2000; Donaldson and Williams, 2018). Autofluorescence in the xylem vascular tissue was visualized by mounting cross-sections in 70 % ethanol (neutral pH) and illuminating them with UV using a Leica DM6B-Z microscope to observe blue auto-fluorescence (340-380 nm excitation and 410-450 nm barrier filters). Images were recorded using a Leica MC190-HD-0518131623 digital camera. These same sections were subsequently mounted in 1N KOH (pH above 10) to observe green auto-fluorescence from ferulic acid using the same settings.

To visualize suberin aliphatics, sections were treated with 5 % Sudan IV, dissolved in 70 % ethanol and illuminated with UV light to produce the typical reddish-brown coloration. These sections were subsequently treated with 1N KOH to detect ferulic acid as described in the previous paragraph. For both ferulic acid and suberin, the HC PL APO or HC PL FLUOTAR objectives of the Leica DM6B-Z microscope were used and images were captured using a Leica MC190-HD-0518131623 color digital camera.

The UV auto-fluorescence signal from xylem vessel walls and surrounding layer of parenchyma cells, tracheids was measured using the LAS X Leica software. Change in ferulate accumulation was quantified from mean fluorescence in the green channels using ImageJ software by selecting area of xylem vessel walls and surrounding layer of parenchyma cells, tracheids showing autoflorescence.

### FT-IR

Dried taproot cross-sections of H7996 and Marmande plants, water-treated or *R. solanacearum-*inoculated by soil soak and containing bacteria 10^5^ CFU g^-1^ taproot tissue were analyzed using a FT-IR spectrophotometer Jasco 4700 with ATR accessory on the range of 300-4000 cm^-1^. The area analyzed was adjacent to the vasculature. All spectra were smoothened to minimize noise and baseline corrected, and the peak due to atmospheric CO_2_ at 2300 cm^-1^ was eliminated for clarity using OriginPro software. Assignation of vibration bands allowed peak identification (Lopes *et al*., 2000; Dorado *et al*., 2001; Martín *et al*., 2005; Lahlali *et al*., 2017), Relative absorbance ratios of peaks of significant importance were calculated by using the absorbance at 1236 cm^-1^ as a reference.

### 2D-NMR

The samples of a pool of 15 tomato plant tap roots, water treated or having a bacterial load of 10^5^ CFU.g^-1^ were milled and extracted sequentially with water (3 x 30 mL), 80% ethanol (3 x 30 mL), and with acetone (2 x 40 mL), by sonicating in an ultrasonic bath during 30 min each time, centrifuging (9000 rpm, 25 min) and eliminating the supernatant. Then, lignin/suberin fraction was enzymatically isolated by hydrolyzing the carbohydrates fraction with Cellulysin (Calbiochem), as previously described (Rico *et al*. 2014). Approximately 20 mg of enzymatic lignin/suberin (ELS) preparation was dissolved in 0.6 mL of DMSO-*d*_6_. Heteronuclear single quantum coherence (HSQC) spectra were acquired on a Bruker AVANCE III 500 MHz spectrometer equipped with a 5 mm TCI cryoprobe, using the experimental conditions previously described (Rico *et al*., 2014). HSQC cross-signals were assigned and quantified as described elsewere (Rencoret *et al*., 2018; del Río *et al*., 2018; Mahmoud *et al*., 2020). In the aromatic region, the correlation signals of G_2_ and S_2,6_ were used to estimate the content of the respective G- and S-lignin units. The C_α_/H_α_ signals of the β–*O*–4′ ethers (A_α_), phenylcoumarans (B_α_), and resinols (C_α_) in the linkages region were used to estimate their relative abundances, whereas the C_γ_/H_γ_ signal was used in the case of cinnamyl alcohol end-units (I_γ_).

### RNA extraction, cDNA synthesis and quantitative RT-PCR analysis

Tomato H7996 and Marmande plants were water-treated or inoculated with *R. solanacearum* by soil soak. Plants with a taproot inoculum of 10^5^ CFU g ^−1^ were selected for RNA extraction. With a sharp razor blade, taproot sections of ∼ 0.5 mm thickness were obtained and the xylem vascular tissues (vascular bundles and surrounding parenchyma cells) were collected and kept in liquid nitrogen. Each sample comprised taproot xylem tissues of 6 plants. RNA was extracted using the Maxwell RSC Plant RNA Kit (Promega) according to the manufactureŕs recommendations. Extracted RNA was treated with RNase-free DNase provided in the kit. cDNA was synthesized from 2 µg RNA using High Capacity cDNA Reverse Transcription Kit (Applied Biosystems, USA). For each reaction 2.5 μl of cDNA (1:20 dilution), 1 μl forward and reverse primer mix (10 μM/μl), 5 μl SYBR Green PCR master mix (Roche) and ultrapure water up to 10 μl was used and analyzed using the LightCycler 480 System (Roche). The amplification program was performed as follows: 10 min at 95°C, followed by 45 cycles of 95°C for 10 sec, 60°C for 30 sec and 72°C for 30 sec. The Elongation Factor 1 alpha houskeeping gene (*eEF1 α, Solyc06g005060*) was used as a reference. All reactions were run in triplicate for each biological replicates. Melting curves and relative quantification of target genes were determined using the software LightCycler V1.5 (Roche). The level of expression relative to the reference gene was calculated using the formula 2^-ΔCT^, where ΔCT = (CT RNA target - CT reference RNA).

### Statistical analysis

Statistical analyses were performed using Statgraphics software. All statistical tests are indicated in the respective figure legends.

## Acknowledgements

The authors would like to thank Gabriel Castrillo (University of Nottingham) and Nico Geldner (University of Lausanne) for inspiring discussions. We also thank Marc Planas-Marquès and all members of the Bacterial plant diseases and cell death lab for helpful comments. We also thank María Pilar López Gresa (IBMCP-UPV) for kindly sharing the tomato THT overexpressor seeds. Research in the lab is funded by the Spanish Ministry of Economy and Competitiveness with grants by the Ministry of Science and Innovation and Innovation State Research Agency PID2019-108595RB-I00/AEI/10.13039/501100011033 (NSC), PID2019-110330GB-C21(MF, OS) through the “Severo Ochoa Programme for Centres of Excellence in R&D” (SEV-2015-0533 and and CEX2019-000902-S), and by the Spanish National Research Council (CISC) pie-201620E081 (JR, AG). AK is the recipient of a Netaji Subhas - Indian Council of Agricultural Research (ICAR) International Fellowship. This work was also supported by the CERCA Programme / Generalitat de Catalunya. We acknowledge support of the publication fee by the CSIC Open Access Publication Support Initiative through its Unit of Information Resources for Research (URICI).

## Author Contribution

AK designed and performed experiments, interpreted data and wrote the manuscript.

MC performed experiments, interpreted data and reviewed the manuscript.

WZ performed experiments.

SS conducted the FT-IR experiments and reviewed the manuscript.

JR isolated the lignin/suberin fractions and conducted the 2D-HSQC NMR analysis, including data interpretation.

AG isolated the lignin/suberin fractions and conducted the 2D-HSQC NMR analysis, including data interpretation.

AL conducted the FT-IR experiments and reviewed the manuscript.

OS conducted histopathology staining experiments, interpreted data and reviewed the manuscript.

MF interpreted data and reviewed the manuscript.

MV designed experiments, interpreted data and review the manuscript.

NSC conceptualized the research, designed experiments, interpreted data and wrote the manuscript.

## Data Availability

The data that support the findings of this study are available from the corresponding author upon reasonable request.

## SUPPLEMENTAL DATA

**Table S1.** Assignments of the correlation signals in the 2D HSQC spectra.

| Label | $\delta_C/\delta_H$ (ppm) | Assignment |
| --- | --- | --- |
| Ty<sub>7</sub> | 34.2/2.62 | C<sub>7</sub>/H<sub>7</sub> in amides of tyramine (**Ty**) |
| Ty<sub>8</sub> | 40.5/3.29 | C<sub>8</sub>/H<sub>8</sub> in amides of tyramine (**Ty**) |
| B<sub>β</sub> | 53.3/3.43 | C<sub>β</sub>/H<sub>β</sub> in phenylcoumarans (**B**) |
| C<sub>β</sub> | 53.5/3.05 | C<sub>β</sub>/H<sub>β</sub> in β-$\beta'$resinols (**C**) |
| MeO | 55.3/3.72 | C/H in aromatic methoxy group |
| A<sub>γ</sub> | 59.7/3.23, 3.58 | C<sub>γ</sub>/H<sub>γ</sub> in β-O-4' alkyl-aryl ethers (**A**) |
| I<sub>γ</sub> | 61.5/4.06 | C<sub>γ</sub>/H<sub>γ</sub> in cinnamyl alcohol end-groups (**I**) |
| B<sub>γ</sub> | 62.6/3.70 | C<sub>γ</sub>/H<sub>γ</sub> in phenylcoumarans (**B**) |
| C<sub>γ</sub> | 71.1/3.80, 4.17 | C<sub>γ</sub>/H<sub>γ</sub> in β-$\beta'$resinols (**B**) |
| A<sub>α</sub> | 71.3/4.79 | C<sub>α</sub>/H<sub>α</sub> in β-O-4' alkyl-aryl ethers (**A**) |
| A<sub>βG</sub> | 83.9/4.27 | C<sub>β</sub>/H<sub>β</sub> in β-O-4' alkyl-aryl ethers (**A**) linked to a G unit |
| C<sub>α</sub> | 84.9/4.67 | C<sub>α</sub>/H<sub>α</sub> in β-$\beta'$resinols (**C**) |
| A<sub>βS</sub> | 83.6/4.28 | C<sub>β</sub>/H<sub>β</sub> in β-O-4' alkyl-aryl ethers (**A**) linked to a S unit |
| B<sub>α</sub> | 86.9/5.45 | C<sub>α</sub>/H<sub>α</sub> in phenylcoumarans (**B**) |
| S<sub>2,6</sub> | 104.0/6.68 | C<sub>2</sub>/H<sub>2</sub> and C<sub>6</sub>/H<sub>6</sub> in syringyl units (**S**) |
| S'<sub>2,6</sub> | 106.3/7.29 | C<sub>2</sub>/H<sub>2</sub> and C<sub>6</sub>/H<sub>6</sub> in Cα-oxidized syringyl units (**S'**) |
| G<sub>2</sub> | 111.1/6.97 | C<sub>2</sub>/H<sub>2</sub> in guaiacyl units (**G**) |
| Ty<sub>3,5</sub> | 114.8/6.64 | C<sub>3</sub>/H<sub>3</sub> and C<sub>5</sub>/H<sub>5</sub> in amides of tyramine (**Ty**) |
| G<sub>5/6</sub> | 114.9/6.79 | C<sub>5</sub>/H<sub>5</sub> and C<sub>6</sub>/H<sub>6</sub> in guaiacyl units (**G**) |
| G<sub>6</sub> | 119.0/6.76 | C<sub>6</sub>/H<sub>6</sub> in guaiacyl units (**G**) |
| I<sub>β</sub> | 128.2/6.21 | C<sub>β</sub>/H<sub>β</sub> in cinnamyl alcohol end-groups (**I**) |
| I<sub>α</sub> | 128.6/6.43 | C<sub>α</sub>/H<sub>α</sub> in cinnamyl alcohol end-groups (**I**) |
| Ty<sub>2,6</sub> | 129.3/6.92 | C<sub>2</sub>/H<sub>2</sub> and C<sub>6</sub>/H<sub>6</sub> in amides of tyramine (**Ty**) |
| U<sub>F</sub> | 129.4/5.31 | -CH=CH- in unsaturated fatty acid structures (**U<sub>F</sub>**) |
| FAm<sub>7</sub> | 138.6/7.31 | C<sub>7</sub>/H<sub>7</sub> in feruloyl amides (**FAm**) |

**Table S2:**
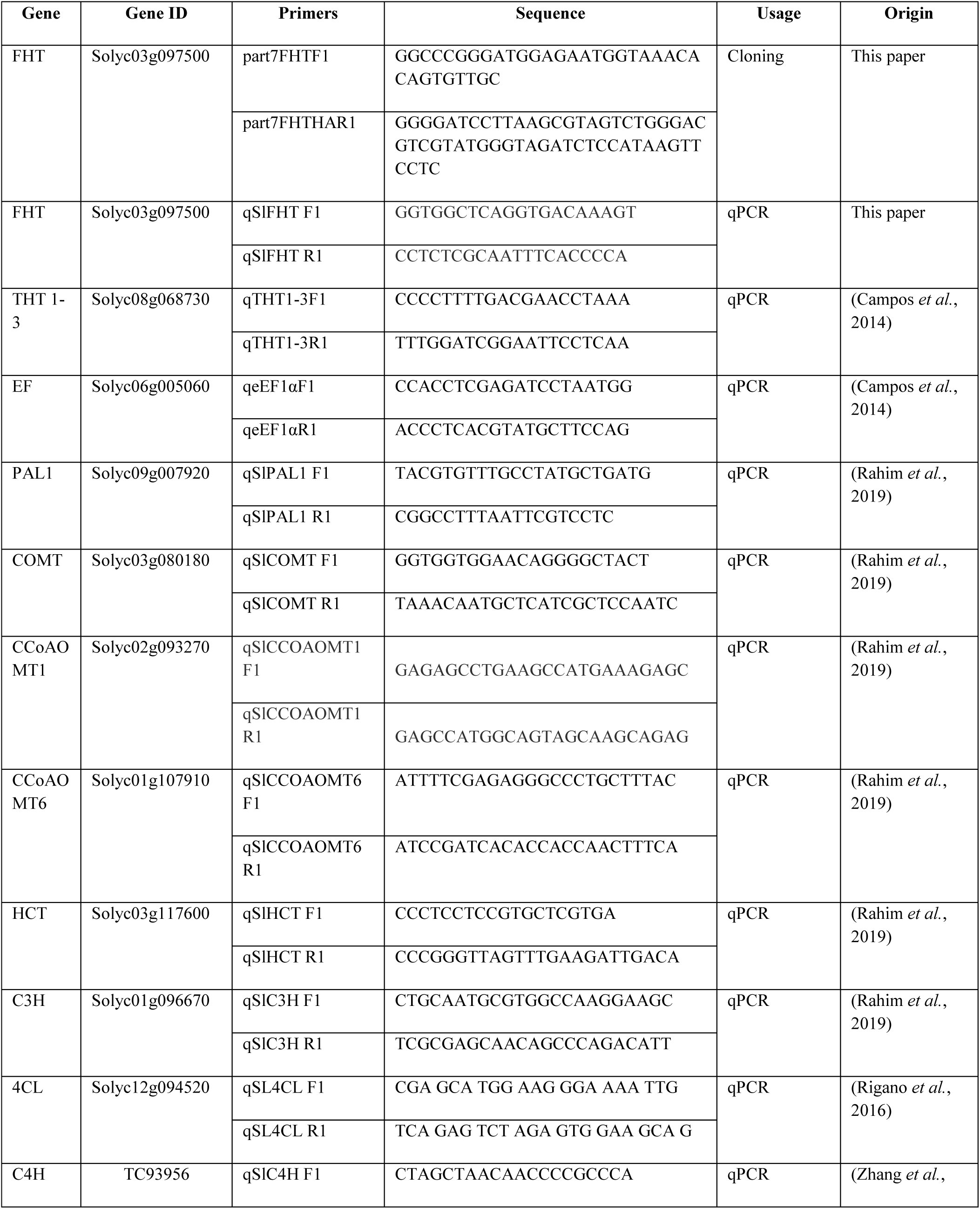
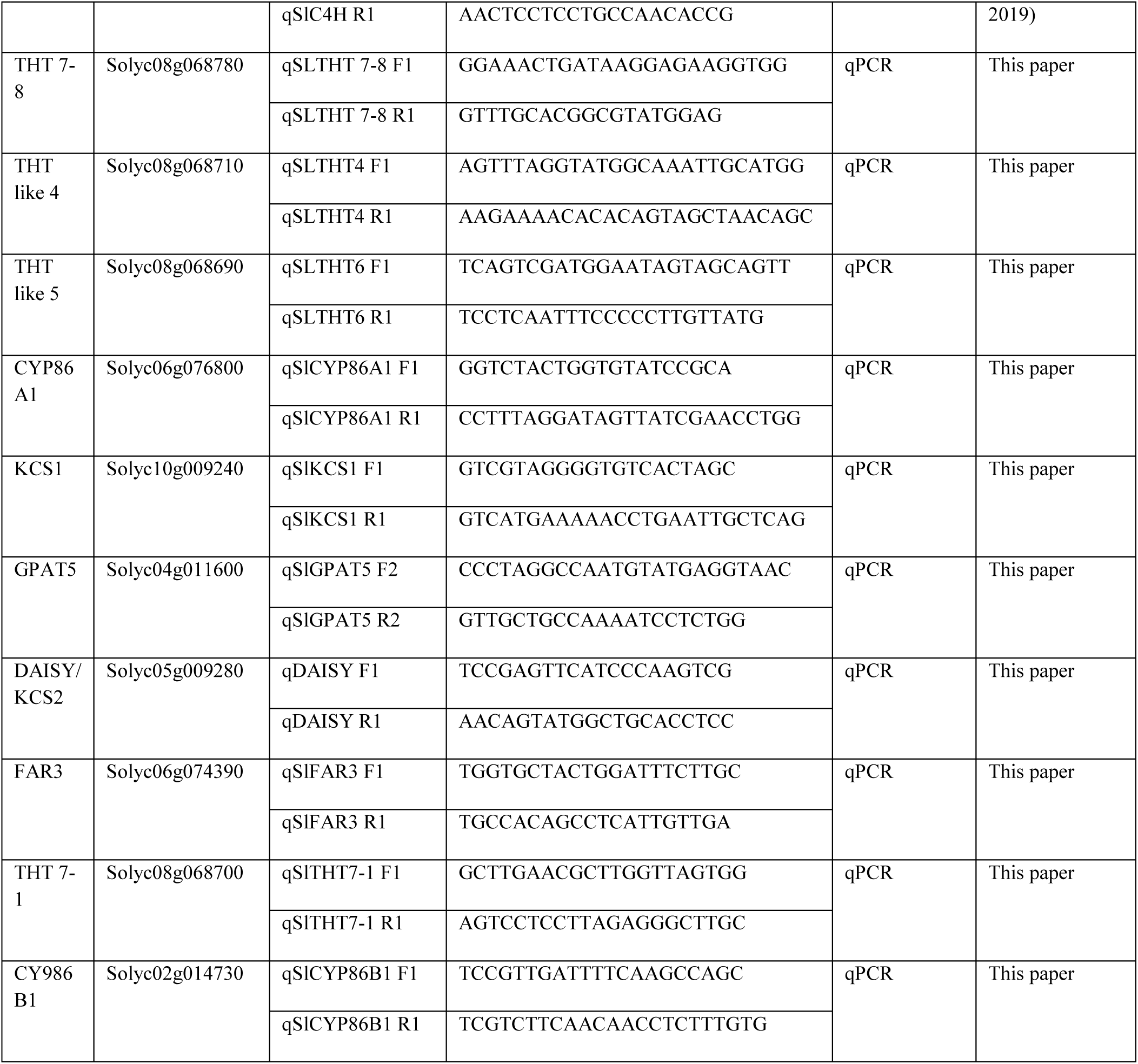
List of primers used for cloning and qPCR analysis.

**Figure S1:**
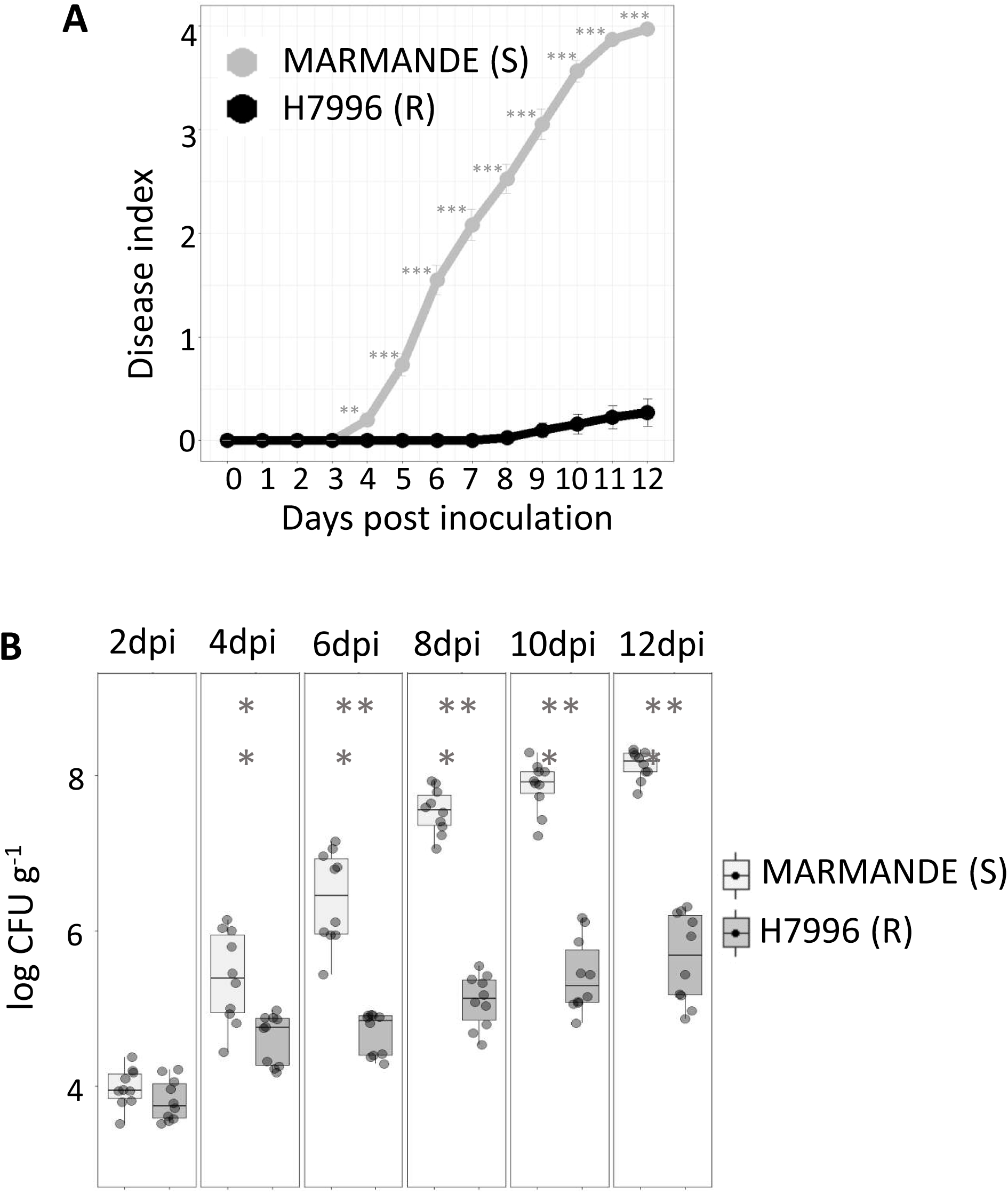
H7996 plants show mild symptoms upon challenge inoculation of *R. solanacearum.* Susceptible (Marmande) and resistant (H7996), 5-week-old tomato plants were inoculated through roots by soil-soak with ∼1×10^7^ CFU/ml of *R. solanacearum* GMI1000 and incubated at 28°C. **(A)** Wilting progress was assayed in both cultivars by rating plants daily on a 0 to 4 disease index scale where 0 = healthy and 4 =100% wilted. Data presented are means ± SE of a representative experiment with n=20 plants for each cultivar, out of a total of 3 experiments. **(B)** The level of in planta colonization by *R. solanacearum* was calculated as colony forming units per gram of fresh taproot tissue (CFU·g ^−1^) at the indicated days post-infection (dpi). Data presented are of a representative experiment with n=10 plants for each time point each cultivar out of a total of 3 experiments. Asterisks indicate statistically significant difference between Marmande and H7996 in a paired Student′s t-test (** p-value of p < 0.01 and *** p-value of p < 0.001). Supports Figure 1 of the main manuscript.

**Figure S2:**
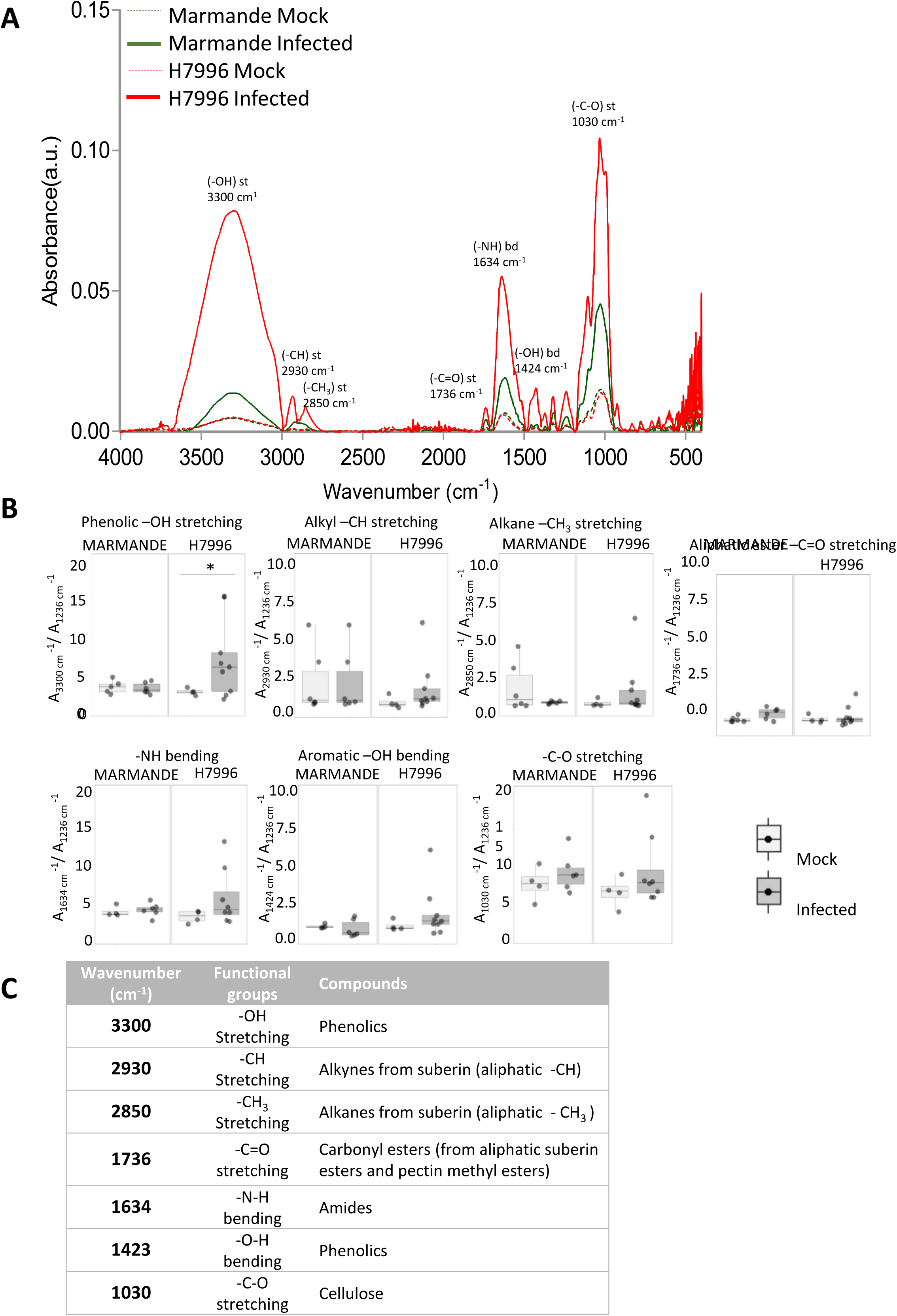
FT-IR showed significantly high induction of phenolics in the xylem vasculature of resistant H7996. Taproot cross-sections of H7996 and Marmande plants, water-treated or *R. solanacearum-*inoculated by soil soak and containing bacteria 10^5^ CFU g^-1^ were analyzed using a FT-IR spectrophotometry in areas adjacent to the vasculature. **(A)** Average absorbance in the range of 500-4000 cm^-1^ is shown for both cultivars water treated or infected with *R. solanacearum.* **(B)** The relative absorbance ratios of the most prominent peaks in (A) were calculated for phenolic –OH stretching (≈ 3300 cm^- 1^), alkyl –OH stretching (≈ 2930 cm^-1^), alkane –CH3 stretching (≈ 2850 cm^-1^), aliphatic ester –C=O stretching (≈ 1736 cm^-1^), – NH bending (≈ 1634 cm^-1^), aromatic –OH bending (≈ 1424 cm^-1^) and -C-O stretching (≈ 1030 cm^-1^) by using the absorbance at 1236 cm^-1^ as a reference. The asterisk (*) indicates statistically significant differences (α=0.05, Student’s t-test). **(C)** Correspondence between wavenumbers (mean of vibrational range), functional groups and compounds. Assignments based on the literature (Dorado *et al*., 2001; Martín *et al*., 2005, 2008; Lahlali *et al*., 2017). Supports Figure 3 of the main manuscript.

**Figure S3:**
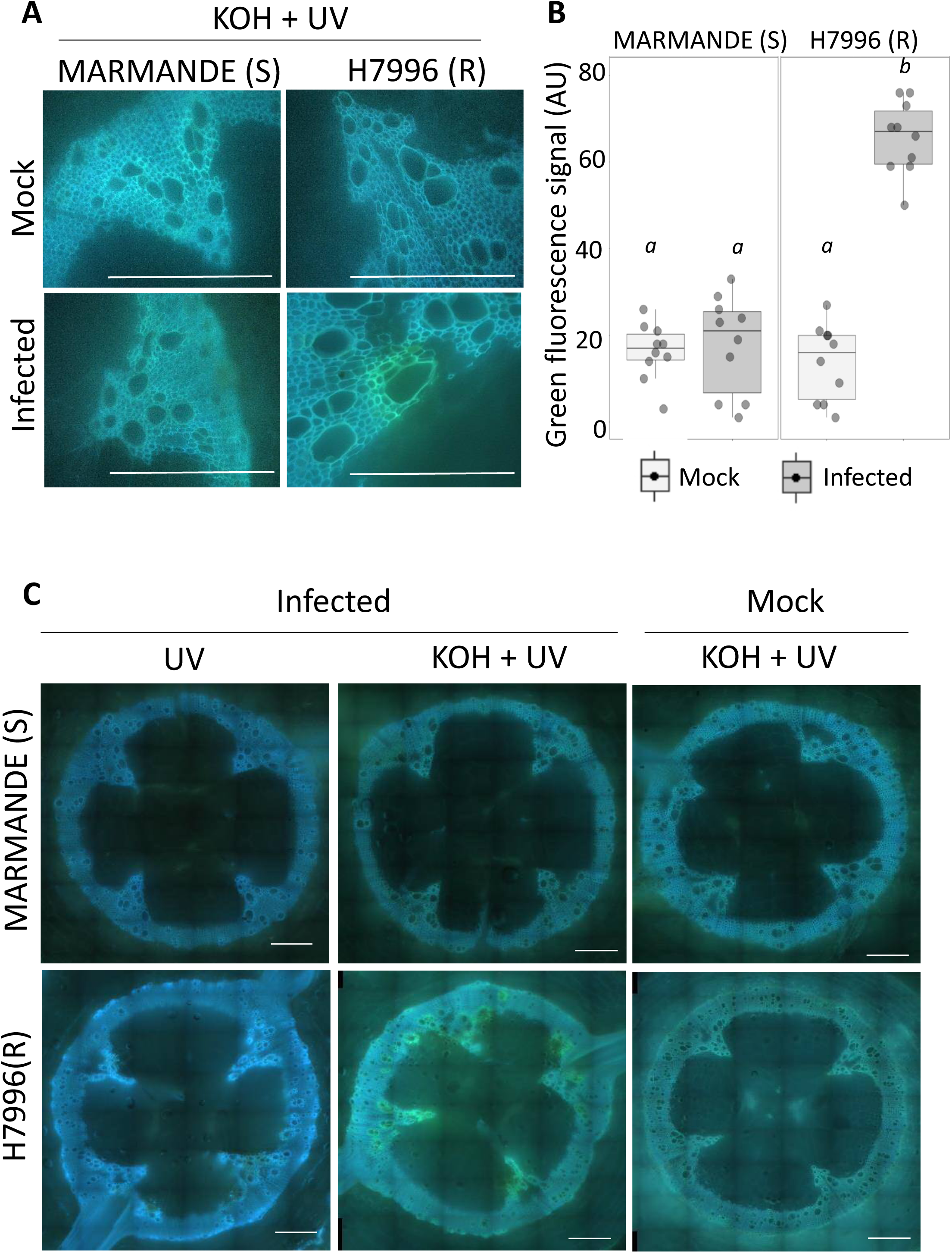
*R. solanacearum-*induced xylem vascular ferulic acid deposition occurs in resistant H7996, but not in susceptible Marmande. Cell wall-bound ferulic acid can be detected by emission of a blue fluorescence with UV excitation at neutral pH, which characteristically changes to stronger green emission under conditions of high pH such as in the presence of alkali. Five-week-old Marmande and H7996 plants were inoculated with ∼1×10^7^ CFU/ml of *R. solanacearum* GMI1000 and incubated at 28°C. Taproot cross-sections were obtained at 9 dpi **(A)** or in plants containing a bacterial load of approximately 10^5^ CFU g ^−1^ of *R. solanacearum* **(B, C)**. **(A)** Autofluorescence emitted from taproot cross-section from mock-treated and infected Marmande and H7996 plants was visualized at 9 dpi under UV before and after treatment with KOH alkali (high pH above 10). In infected H7996 a green/turquoise color appears in vessel walls and surrounding xylem parenchyma cells, indicative of ferulic acid deposition in the cell walls. Scale bars = 500 µm. **(B)** The same as in (A) but cross-sections were obtained from plants containing a bacterial load of approximately 10^5^ CFU g ^−1^ of *R. solanacearum.* Scale bars = 500 µm. **(C)** Green fluorescence from ferulate deposits in the xylem and surrounding parenchyma cells was measured using ImageJ. Box-and-whisker plots show data from a single representative experiment (n =6) out of a total of 3. Different letters indicate statistically significant differences (α=0.05, Fisher’s least significant difference test). Supports Figure 4 of the main manuscript.

**Figure S4:**
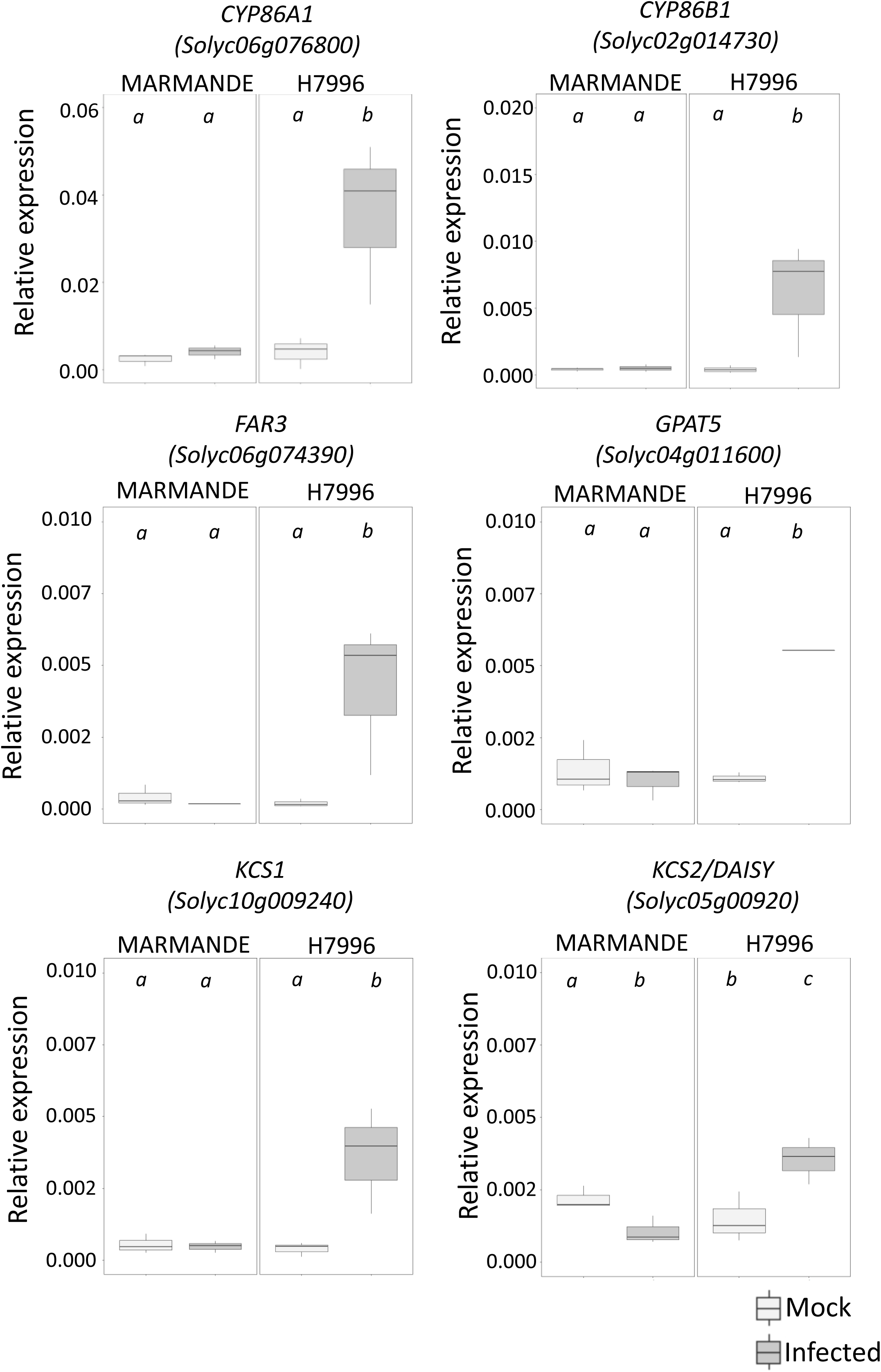
Expression of suberin biosynthetic genes in xylem vasculature of taproots upon infection of *R. solanacearum*. Expression levels of tomato putative orthologs of the suberin fatty acid pathway were analyzed by qPCR in H7996 and Marmande plants infected with *R. solanacearum* or mock-treated. Xylem vascular tissue comprising of metaxylems and surrounding parenchyma cells was collected from infected plants with a *R. solanacearum* inoculum of 10^5^ CFU g^−1^ in the taproot or mock-inoculated plants of a similar age. Relative expression values were calculated using the Elongation Factor 1 alpha (eEF1 α) gene as reference. Three biological replicates (n=3) were used, and taproots of 6 plants were used in each replicate. Different letters indicate statistically significant differences (α=0.05, Fisher’s least significant difference test). Supports Figure 5 of the main manuscript.

**Figure S5:**
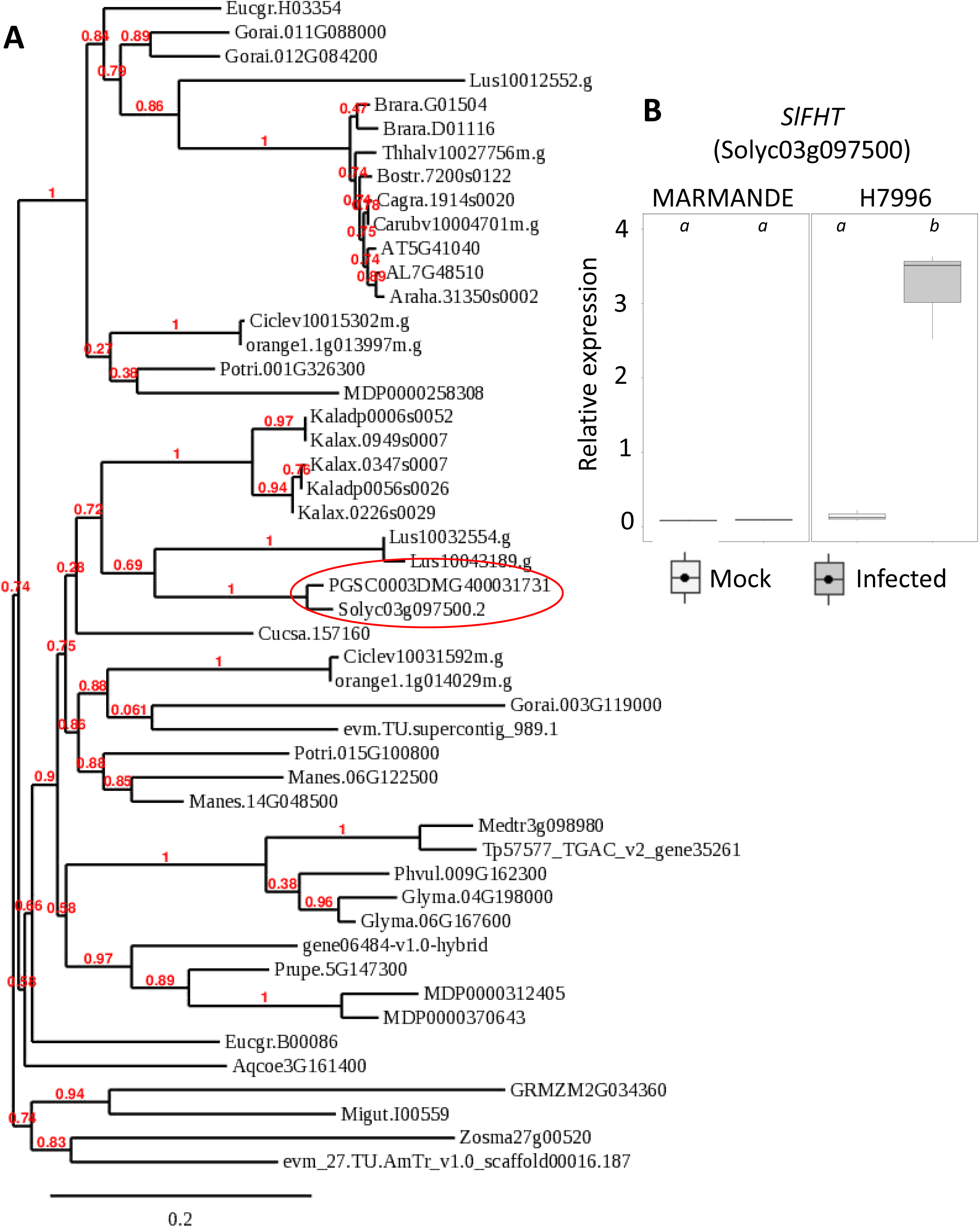
Phylogeny of Feruloyl transferase (FHT) orthologues in different plant species and expression of the putative tomato FHT ortholog in response to *Ralstonia solanacearum* infection. **(A)** Protein homologs of potato *FHT* gene (*PGSC0003DMG400031731*) were obtained from www.phytozome.jgi.doe.gov and matches with more than 80 % similarity were used for phylogenetic analysis using www.phylogeny.fr. **(B)** Gene expression of the putative tomato *FHT* ortholog (*Solyc03g097500*) was analyzed by qPCR. Relative expression levels were calculated using the Elongation Factor 1 alpha (*eEF1 α, Solyc06g005060*) as the reference gene. H7996 and Marmande plants, containing a *R. solanacearum* inoculum of 10^5^ CFU g^−1^ in the taproot were selected. Xylem vascular tissue, comprising of metaxylems and surrounding parenchyma cells was collected from taproots for RNA extraction and cDNA synthesis. Similarly, xylem tissue was collected from Marmande mock plants and H7996 mock plants. Three biological replicates (n=3) were used, and taproots of 6 plants were used in each replicate. Different letters indicate statistically significant differences (α=0.05, Fisher’s least significant difference test). Supports Figure 5 of the main manuscript.

**Figure S6:**
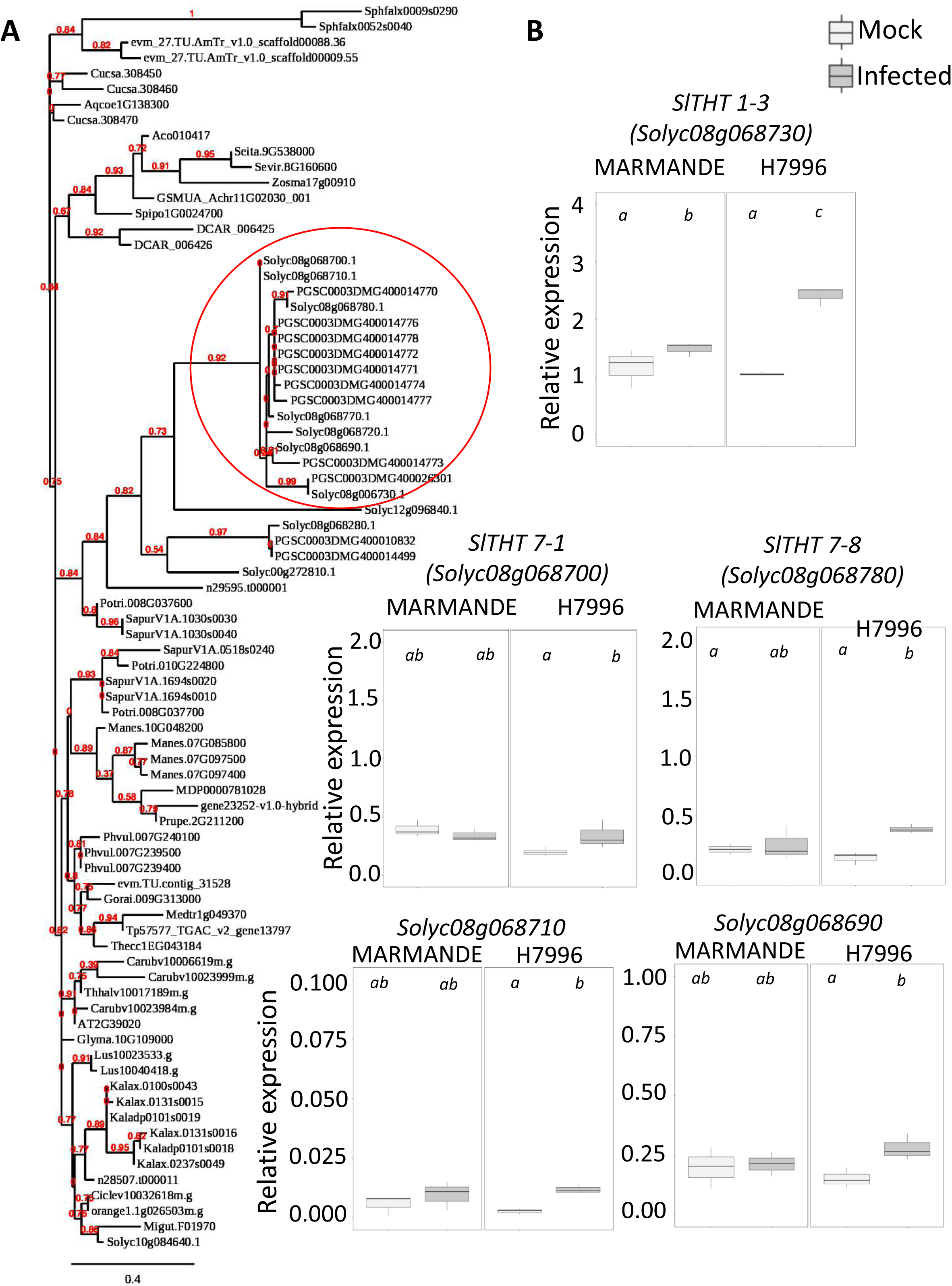
Phylogeny of tyramine hydroxycinnamoyl transferase (THT) orthologues in different plant species and expression of the tomato THT gene family members in response to *R. solanacearum* infection. **(A)** Protein homologs of tomato *THT1-3* gene (*Solyc08g068730*) were obtained from www.phytozome.jgi.doe.gov and matches with more than 60 % similarity were used for phylogenetic analysis using the webpage www.phylogeny.fr. **(B)** Gene expression of the tomato *THT* gene family members was analyzed by qPCR. Relative expression levels were calculated using the Elongation Factor 1 alpha (*eEF1 α, Solyc06g005060*) as the reference gene. H7996 and Marmande plants, containing a *R. solanacearum* inoculum of 10^5^ CFU g^−1^ in the taproot were selected. Xylem vascular tissue, comprising of metaxylems and surrounding parenchyma cells was collected from taproots for RNA extraction and cDNA synthesis. Similarly, xylem tissue was collected from Marmande mock plants and H7996 mock plants. Three biological replicates (n=3) were used, and taproots of 6 plants were used in each replicate. Different letters indicate statistically significant differences (α=0.05, Fisher’s least significant difference test). Supports Figure 5 of the main manuscript.

**Figure S7:**
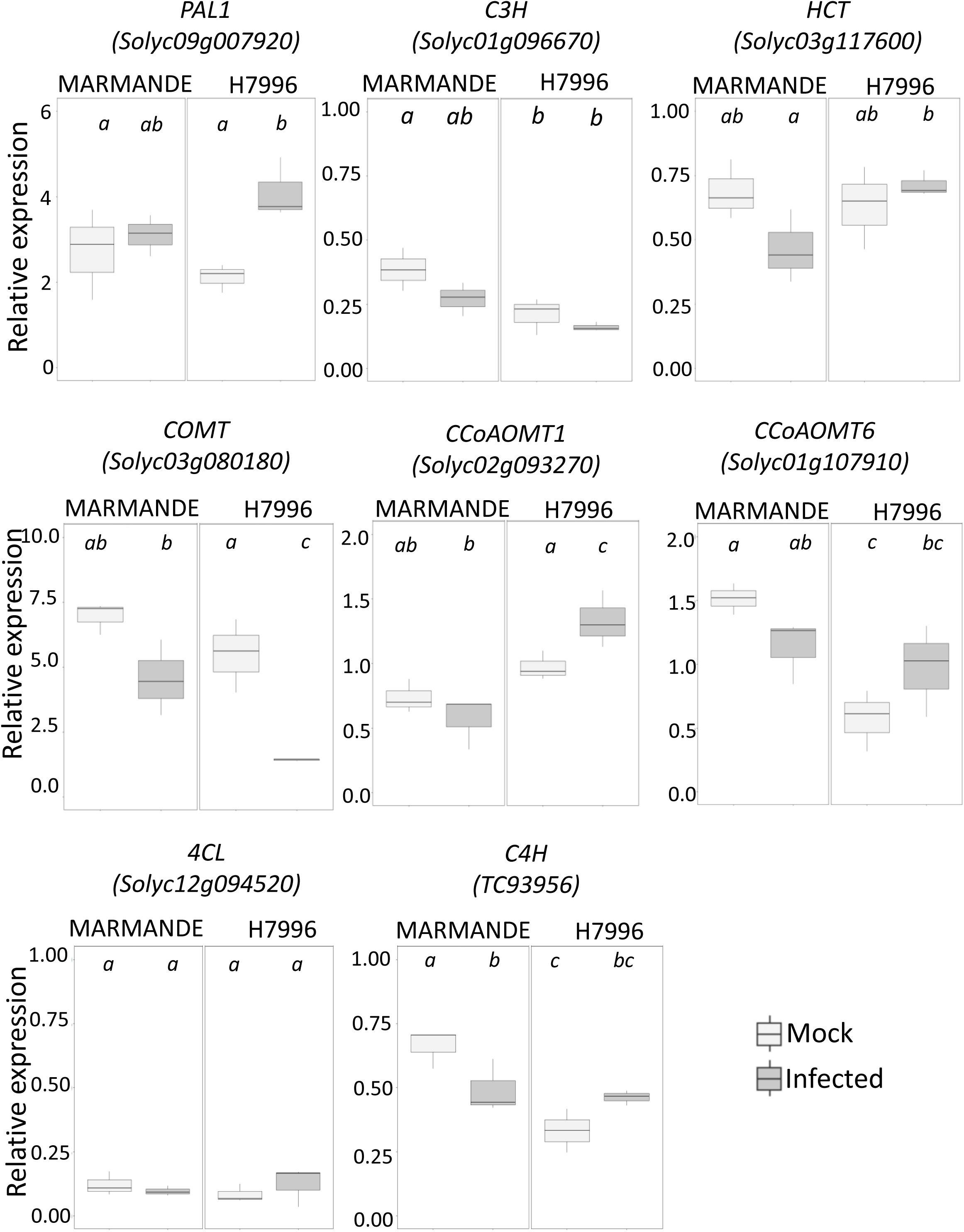
Expression of phenylpropanoid pathway genes in xylem vasculature of taproots upon invasion of *R. solanacearum*. Expression levels of tomato putative orthologs of the phenylpropanoid pathway were analyzed by qPCR in H7996 and Marmande plants infected with *R. solanacearum* or mock-treated. Xylem vascular tissue comprising of metaxylems and surrounding parenchyma cells was collected from infected plants with a *R. solanacearum* inoculum of 10^5^ CFU g^−1^ in the taproot or mock-inoculated plants of a similar age. Relative expression values were calculated using the Elongation Factor 1 alpha (*eEF1 α*) gene as reference. Three biological replicates (n=3) were used, and taproots of 6 plants were used in each replicate. Different letters indicate statistically significant differences (α=0.05, Fisher’s least significant difference test). Supports Figure 5 of the main manuscript.

**Figure S8:**
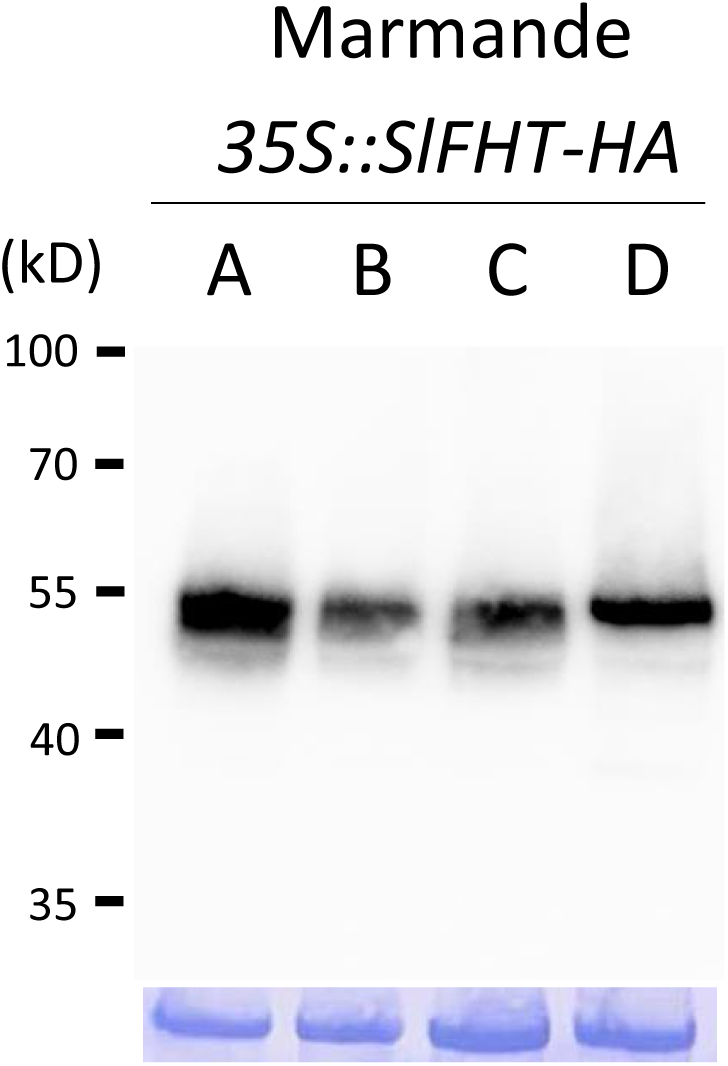
Immunoblot of SlFHT-HA in independent Marmande tomato lines expressing *35S::SlFHT-HA* (Marmande). Immunoblot using anti-HA antibody showing SlFHT-HA protein (predicted protein size: 49kDa) levels of 4 independent transgenic lines (A, B, C D) stably overexpressing *SlFHT-HA* on a susceptible Marmande background (*35S::SlFHT-HA*). In the bottom panel Coomasie blue staining showing similar protein load in all the lanes is shown. Supports Figure 6 of the main manuscript.

**Figure S9:**
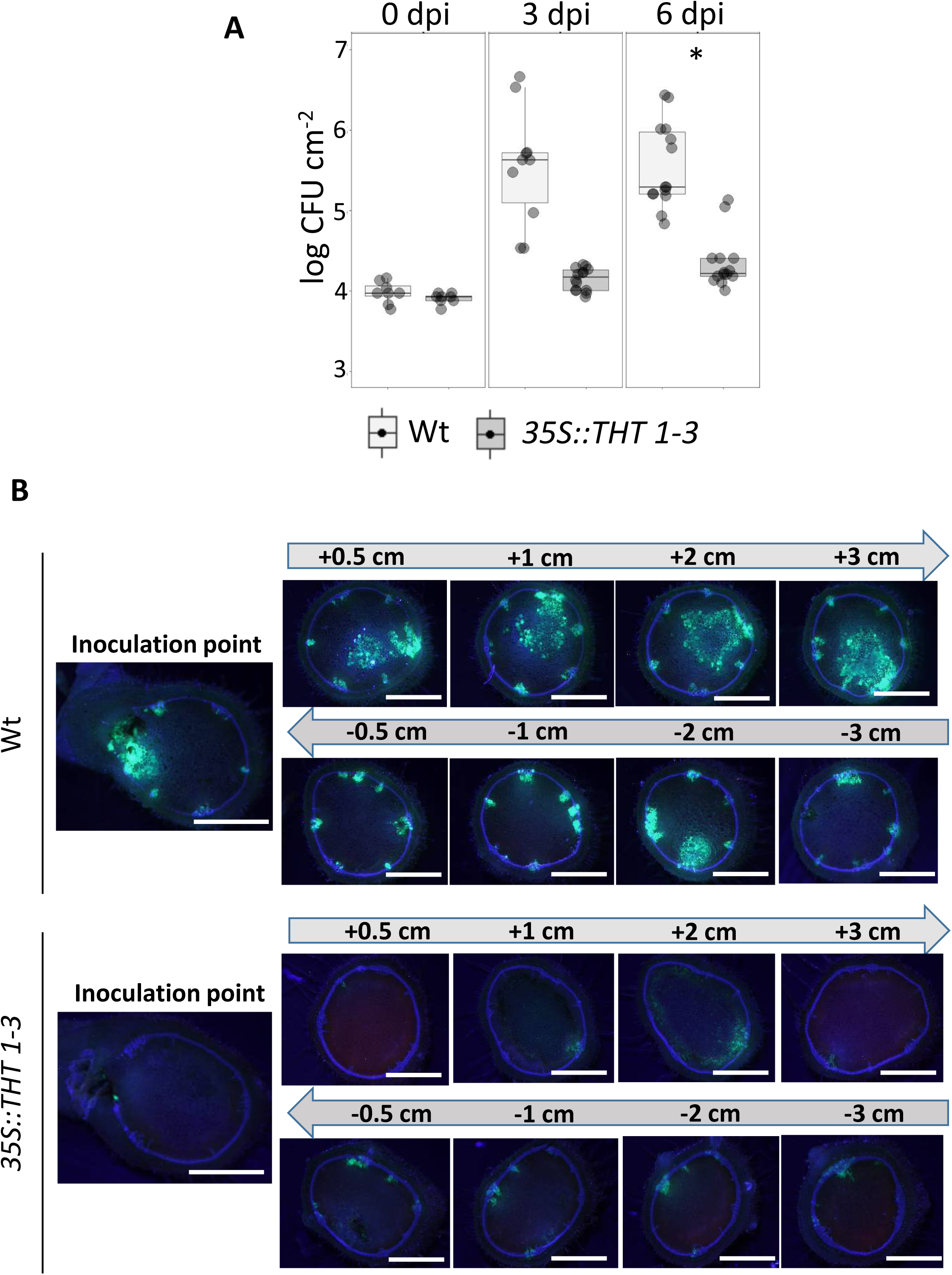
Overexpression of *SlTHT1-3* in tomato results in restricted colonization by *R. solanacearum*. **(A)** Growth of *R. solanacearum* GMI1000 in planta was monitored in leaves of *35S::THT1-3* transgenic plant compared with Wt Moneymaker tomato lines over time. The bacterium was vacuum infiltrated into the leaves at a concentration of ∼1×10^5^ CFU/ml and growth was recorded at 0, 3 and 6 dpi. Box-and-whisker plots show data of 6 to 8 independent plants (n=6-8) from a representative experiment out of 3. Asterisk indicates statistically significant difference between Wt and overexpression line in a paired Student′s t-test (* corresponds to p-value of p < 0.05). **(B)** Representative images of tomato stem cross-sections showing colonization by the *R. solanacearum* GMI1000 GFP reporter strain at 6 dpi. *R. solanacearum* was directly injected into the xylem vasculature of the first internode through the petiole at a concentration of 10^5^ CFU ml ^−1^. Colonization progress was analyzed at the point of inoculation, at higher (+0.5, +1, +2 and +3 cm) and lower -0.5, -1, -2 and - 3 cm) sections. Images from a representative experiment out of 3 with *n*=5 plants each. Scale bar = 2 mm. Supports Figure 7 of the main manuscript.

## Parsed Citations

Álvarez, B., Biosca, E.G., and López, M.M. (2010). On the life of Ralstonia solanacearum, a destructive bacterial plant pathogen. In Technology and education topics in applied microbiology and microbial biotechnology, A. Méndez-Vilas, ed (Badajoz: Formatex), pp. 267–279.

Andersen, T.G., Molina, D., Kilian, J., Franke, R.B., Ragni, L., and Geldner, N. (2021). Tissue-autonomous phenylpropanoid production is essential for establishment of root barriers. Curr. Biol. 31: 965–977.

Araujo, L., Bispo, W.M.S., Cacique, I.S., Moreira, W.R., and Rodrigues, F.A. (2014). Resistance in mango against infection by Ceratocystis fimbriata. Phytopathology 104: 820–833.

De Ascensao, A.R.D.C.F. and Dubery, I.A. (2000). Panama disease: cell wall reinforcement in banana roots in response to elicitors from Fusariumoxysporum f. sp. cubense race four. Phytopathology 90: 1173–1180.

Baayen, R.P. and Elgersma, D.M. (1985). Colonization and histopathology of susceptible and resistant carnation cultivars infected with Fusariumoxysporum f. sp. dianthi. Netherlands J. Plant Pathol. 91: 119–135.

Benhamou, N. (1995). Ultrastructural and cytochemical aspects of the response of eggplant parenchyma cells in direct contact with Verticillium-infected xylem vessels. Physiol. Mol. Plant Pathol. 46: 321–338.

Bernards, M., Lopez, M., Zajicek, J., and Lewis, N. (1995). Hydroxycinnamic acid-derived polymers constitute the polyaromatic domain of suberin. J. Biol. Chem. 270: 7382–7386.

Bernards, M.A. (2002). Demystifying suberin. Can. J. Bot. 80: 227–240.

Bernards, M.A. and Lewis, N.G. (1998). The macromolecular aromatic domain in suberized tissue: a changing paradigm. Phytochemistry 47: 915–933.

Biggs, A. (1984). Intracellular suberin: occurrence and detection in tree bark. IAWA Bull. 5: 243–248.

Campos, L., Lisón, P., López-Gresa, M.P., Rodrigo, I., Zacarés, L., Conejero, V., and Bellés, J.M. (2014). Transgenic tomato plants overexpressing tyramine N -hydroxycinnamoyltransferase exhibit elevated hydroxycinnamic acid amide levels and enhanced resistance to Pseudomonas syringae. Mol. Plant-Microbe Interact. 27: 1159–1169.

Carnachan, S.M. and Harris, P.J. (2000). Ferulic acid is bound to the primary cell walls of all gymnosperm families. Biochem. Syst. Ecol. 28: 865–879.

Correia, V.G., Bento, A., Pais, J., Rodrigues, R., Haliński, P., Frydrych, M., Greenhalgh, A., Stepnowski, P., Vollrath, F., King, A.W.T., and Pereira, C.S. (2020). The molecular structure and multifunctionality of the cryptic plant polymer suberin. Mater. Today Bio 5: 100039.

Cruz, A.P.Z., Ferreira, V., Pianzzola, M.J., Siri, M.I., Coll, N.S., and Valls, M. (2014). Anovel, sensitive method to evaluate potato germplasm for bacterial wilt resistance using a luminescent Ralstonia solanacearumreporter strain. Mol. Plant-Microbe Interact. 27: 277–285.

Digonnet, C., Martinez, Y., Denancé, N., Chasseray, M., Dabos, P., Ranocha, P., Marco, Y., Jauneau, A., and Goffner, D. (2012). Deciphering the route of Ralstonia solanacearumcolonization in Arabidopsis thaliana roots during a compatible interaction: Focus at the plant cell wall. Planta 236: 1419–1431.

Donaldson, L. (2020). Autofluorescence in plants. Molecules 25: 2393.

Donaldson, L. and Williams, N. (2018). Imaging and spectroscopy of natural fluorophores in pine needles. Plants 7: 10.

Dorado, J., Almendros, G., Field, J.A., and Sierra-alvarez, R. (2001). Infrared spectroscopy analysis of hemp (Cannabis sativa) after selective delignification by Bjerkandera sp. at different nitrogen levels. Enzyme Microb. Technol. 28: 550–559.

Falter, C., Ellinger, D., Von Hulsen, B., Heim, R., and Voigt, C.A. (2015). Simple preparation of plant epidermal tissue for laser microdissection and downstreamquantitative proteome and carbohydrate analysis. Front. Plant Sci. 6: 194.

Faragher, J.D. and Brohier, R.L. (1984). Anthocyanin accumulation in apple skin during ripening: regulation by ethylene and phenylalanine ammonia-lyase. Sci. Hortic. 22: 89–96.

Fattorusso, E., Lanzotti, V., and Taglialatela-Scafati, O. (1999). Antifungal N-feruloylamides fromroots of two alliumspecies. Plant Biosyst. 133: 199–203.

Figueiredo, R., Portilla Llerena, J.P., Kiyota, E., Ferreira, S.S., Cardeli, B.R., de Souza, S.C.R., dos Santos Brito, M., Sodek, L., Cesarino, I., and Mazzafera, P. (2020). The sugarcane ShMYB78 transcription factor activates suberin biosynthesis in Nicotiana benthamiana. Plant Mol. Biol. 104: 411–427.

Gleave, A.P. (1992). Aversatile binary vector system with a T-DNAorganisational structure conducive to efficient integration of cloned DNAinto the plant genome. Plant Mol. Biol. 20: 1203–1207.

Gou, J.-Y., Yu, X.-H., and Liu, C.-J. (2009). Ahydroxycinnamoyltransferase responsible for synthesizing suberin aromatics in Arabidopsis. Proc. Natl. Acad. Sci. 106: 18855–18860.

Graça, J. (2010). Hydroxycinnamates in suberin formation. Phytochem. Rev. 9: 85–91.

Graça, J. (2015). Suberin: the biopolyester at the frontier of plants. Front. Chem. 3: 62.

Grimault, V., Anais, G., and Prior, P. (1994). Distribution of Pseudomonas solanacearumin the stemtissues of tomato plants with different levels of resistance to bacterial wilt. Plant Pathol. 43: 663–668.

Hao, Z. et al. (2014). Loss of arabidopsis GAUT12/IRX8 causes anther indehiscence and leads to reduced G lignin associated with altered matrix polysaccharide deposition. Front. Plant Sci. 5: 357.

Harris, P.J. and Trethewey, J.A.K. (2010). The distribution of ester-linked ferulic acid in the cell walls of angiosperms. Phytochem. Rev. 9: 19–33.

He, M. and Ding, N. (2020). Plant unsaturated fatty acids : multiple roles in stress response. Front. Plant Sci. 11: 562785.

Howles, P.A., Sewalt, V.J.H., Paiva, N.L., Elkind, Y., Bate, N.J., Lamb, C., and Dixon, R.A. (1996). Overexpression of L-phenylalanine ammonia-lyase in transgenic tobacco plants reveals control points for flux into phenylpropanoid biosynthesis. Plant Physiol. 112: 1617– 1624.

Iiyama, K., Lam, T.B.T., and Stone, B. (2020). Covalent cross-links in the cell wall. Plant Physiol. 104: 315–320.

Ishihara, T., Mitsuhara, I., Takahashi, H., and Nakaho, K. (2012). Transcriptome analysis of quantitative resistance-specific response upon Ralstonia solanacearuminfection in tomato. PLoS One 7(10): e46763.

Jones, J.D.G. and Dangl, J.L. (2006). The plant immune system. Nature 444: 323–329.

Kashyap, A., Planas-marquès, M., Capellades, M., Valls, M, and Coll, N.S. (2021). Blocking intruders : inducible physico-chemical barriers against plant vascular wilt pathogens. J. Exp. Bot. 72: 184–198.

Kim, S.G., Hur, O.S., Ro, N.Y., Ko, H.C., Rhee, J.H., Sung, J.S., Ryu, K.Y., Lee, S.Y., and Baek, H.J. (2016). Evaluation of resistance to Ralstonia solanacearumin tomato genetic resources at seedling stage. Plant Pathol. J. 32: 58–64.

Kutscha, N.P. and Gray, J.R. (1972). The suitability of certain stains for studying lignification in balsamfir, Abies balsamina (L.) Mill. Tech. Bull. 53: 1–51.

Lahlali, R., Song, T., Chu, M., Yu, F., Kumar, S., Karunakaran, C., and Peng, G. (2017). Evaluating changes in cell-wall components associated with clubroot resistance using fourier transforminfrared spectroscopy and RT-PCR. International J. Mol. Sci. 18: 2058.

Lashbrooke, J., Cohen, H., Levy-Samocha, D., Tzfadia, O., Panizel, I., Zeisler, V., Massalha, H., Stern, A., Trainotti, L., Schreiber, L., Costa, F., and Aharoni, A. (2016). MYB107 and MYB9 homologs regulate suberin deposition in angiosperms. Plant Cell 28: 2097–2116.

Legay, S., Guerriero, G., André, C., Guignard, C., Cocco, E., Charton, S., Boutry, M., Rowland, O., and Hausman, J.F. (2016). MdMyb93 is a regulator of suberin deposition in russeted apple fruit skins. New Phytol. 212: 977–991.

Liu, H., Zhang, S., Schell, M.A., and Denny, T.P. (2005). Pyramiding unmarked deletions in Ralstonia solanacearumshows that secreted proteins in addition to plant cell-wall-degrading enzymes contribute to virulence. Mol. Plant-Microbe Interact. 18: 1296–1305.

Lopes, M. H., Neto, C. P., Barros, A. S., Rutledge, D., Delgadillo, I., and Gil, A. M. (2000). Quantitation of aliphatic suberin in Quercus suber L. cork by FTIR spectroscopy and solid-state 13C-NMR spectroscopy. Biopolymers 57: 344–351.

Lowe-Power, T.M., Khokhani, D., and Allen, C. (2018). How Ralstonia solanacearumexploits and thrives in the flowing plant xylem environment. Trends Microbiol. 26: 929–942.

Lulai, E.C. and Corsini, D.L. (1998). Differential deposition of suberin phenolic and aliphatic domains and their roles in resistance to infection during potato tuber (Solanumtuberosum L.) wound-healing. Physiol. Mol. Plant Pathol. 53: 209–222.

Macoy, D.M., Kim, W.Y., Lee, S.Y., and Kim, M.G. (2015). Biotic stress related functions of hydroxycinnamic acid amide in plants. J. Plant Biol. 58: 156–163.

Mahmoud, A.B., Danton, O., Kaiser, M., Han, S., Moreno, A., Algaffar, S.A., Khalid, S., Oh, W.K., Hamburger, M., and Mäser, P. (2020). Lignans, amides, and saponins from Haplophyllum tuberculatumand their antiprotozoal activity. Molecules 25: 2825.

Mangin, B., Thoquet, P., Olivier, J., and Grimsley, N.H. (1999). Temporal and multiple quantitative trait loci analyses of resistance to bacterial wilt in tomato permit the resolution of linked loci. Genetics 151: 1165–1172.

Martín, J.A., Solla, A., Coimbra, M.A., and Gil, L. (2005). Metabolic distinction of Ulmus minor xylem tissues after inoculation with Ophiostoma novo-ulmi. Phytochemistry 66: 2458–2467.

Martín, J.A., Solla, A., Domingues, M.R., Coimbra, M.A., and Gil, L. (2008). Exogenous phenol increase resistance of Ulmus minor to dutch elmdisease through formation of suberin-like compounds on xylem tissues. Environ. Exp. Bot. 64: 97–104.

Mazier, M., Flamain, F., Nicolaï, M., Sarnette, V., and Caranta, C. (2011). Knock-down of both eIF4E1 and eIF4E2 genes confers broad-spectrumresistance against potyviruses in tomato. PLoS One 6(12): e29595.

Mnich, E. et al. (2020). Phenolic cross-links: building and de-constructing the plant cell wall Ewelina. Nat. Prod. Rep. 37: 919–961.

Molina, I., Li-Beisson, Y., Beisson, F., Ohlrogge, J.B., and Pollard, M. (2009). Identification of an Arabidopsis feruloyl-coenzyme a transferase required for suberin synthesis. Plant Physiol. 151: 1317–1328.

Nakaho, K., Hibino, H., and Miyagawa, H. (2000). Possible mechanisms limiting movement of Ralstonia solanacearumin resistant tomato tissues. J. Phytopathol. 148: 181–190.

Nakaho, K., Inoue, H., Takayama, T., and Miyagawa, H. (2004). Distribution and multiplication of Ralstonia solanacearumin tomato plants with resistance derived fromdifferent origins. J. Gen. Plant Pathol. 70:115–119.

Negrel, J., Javelle, F., and Paynot, M. (1993). Wound-induced tyramine hydroxycinnamoyl transferase in Potato (Solanumtuberosum) tuber discs. J. Plant Physiol. 142: 518–524.

Negrel, J., Pollet, B., and Lapierre, C. (1996). Ether-linked ferulic acid amides in natural and wound periderms of potato tuber. Phytochemistry 43: 1195–1199.

Novaes, E., Kirst, M., Chiang, V., Winter-sederoff, H., and Sederoff, R. (2010). Lignin and biomass: A negative correlation for wood formation and lignin content in trees. Plant Physiol. 154: 555–561.

Novo, M., Silvar, C., Merino, F., Martínez-Cortés, T., Lu, F., Ralph, J., and Pomar, F. (2017). Deciphering the role of the phenylpropanoid metabolism in the tolerance of Capsicumannuum L. to Verticilliumdahliae Kleb. Plant Sci. 258: 12–20.

Pérez-Donoso, A.G., Sun, Q., Caroline Roper, M., Carl Greve, L., Kirkpatrick, B., and Labavitch, J.M. (2010). Cell wall-degrading enzymes enlarge the pore size of intervessel pit membranes in healthy and Xylella fastidiosa-infected grapevines. Plant Physiol. 152: 1748–1759.

Planas-Marquès, M., Bernardo-Faura, M., Paulus, J., Kaschani, F., Kaiser, M., Valls, M., Van Der Hoorn, R.A.L., and Coll, N.S. (2018). Protease activities triggered by Ralstonia solanacearuminfection in susceptible and tolerant tomato lines. Mol. Cell. Proteomics 17: 1112–1125.

Planas-Marquès, M., Kressin, J.P., Kashyap, A., Panthee, D.R., Louws, F.J., Coll, N.S., and Valls, M. (2019). Four bottlenecks restrict colonization and invasion by the pathogen Ralstonia solanacearumin resistant tomato. J. Exp. Bot. 71: 2157–2171.

Pomar, F., Merino, F., and Barceló, A.R. (2002). O-4-linked coniferyl and sinapyl aldehydes in lignifying cell walls are the main targets of the Wiesner (phloroglucinol-HCl) reaction. Protoplasma 220: 17–28.

Pomar, F., Novo, M., Bernal, M.A., Merino, F., Barceló, A.R., and Barceló, A.R. (2004). Changes in stemlignins (monomer composition and crosslinking) and peroxidase are related with the maintenance of leaf photosynthetic integrity during Verticilliumwilt in Capsicum annuum. New Phytol. 163: 111–123.

Potter, C., Harwood, T., Knight, J., and Tomlinson, I. (2011). Learning fromhistory, predicting the future: The UK dutch elmdisease outbreak in relation to contemporary tree disease threats. Philos. Trans. R. Soc. B Biol. Sci. 366: 1966–1974.

Pouzoulet, J., Jacques, A., Besson, X., Dayde, J., and Mailhac, N. (2013). Histopathological study of response of Vitis vinifera cv. Cabernet Sauvignon to bark and wood injury with and without inoculation by Phaeomoniella chlamydospora. Phytopathol. Mediterr. 52: 313–323.

Pradhan Mitra, P. and Loqué, D. (2014). Histochemical staining of Arabidopsis thaliana secondary cell wall elements. J. Vis. Exp.: 87: e51381.

Ralph, J. and Landucci, L. (2010). NMR of lignins. In Lignin and lignans: advances in chemistry, J.A. Heitner, C., Dimmel, D. R., Schmidt, ed, pp. 137–243.

Razem, F.A. and Bernards, M.A. (2002). Hydrogen peroxide is required for poly(phenolic) domain formation during wound-induced suberization. J. Agric. Food Chem. 50: 1009–1015.

Rencoret, J., Kim, H., Evaristo, A.B., Gutiérrez, A., Ralph, J., and Del Río, J.C. (2018). Variability in lignin composition and structure in cell walls of different parts of macaúba (Acrocomia aculeata) Palm Fruit. J. Agric. Food Chem. 66: 138–153.

Rico, A., Rencoret, J., Del Río, J.C., Martínez, A.T., and Gutiérrez, A. (2014). Pretreatment with laccase and a phenolic mediator degrades lignin and enhances saccharification of Eucalyptus feedstock. Biotechnol. Biofuels 7: 6.

del Río, J.C., Rencoret, J., Gutiérrez, A., Kim, H., and Ralph, J. (2018). Structural characterization of lignin from Maize (Zea mays L.) fibers: evidence for diferuloylputrescine incorporated into the lignin polymer in Maize kernels. J. Agric. Food Chem. 66: 4402–4413.

Rioux, D., Blais, M., Nadeau-Thibodeau, N., Lagacé, M., Des Rochers, P., Klimaszewska, K., and Bernier, L. (2018). First extensive microscopic study of butternut defense mechanisms following inoculation with the canker pathogen Ophiognomonia clavigignenti-juglandacearumreveals compartmentalization of tissue damage. Phytopathology 108: 1237–1252.

Rioux, D., Nicole, M., Simard, M., and Ouellette, G.B. (1998). Immunocytochemical evidence that secretion of pectin occurs during gel (gum) and tylosis formation in trees. Phytopathology 88: 494–505.

Rittinger, P.A., Biggs, A.R., and Peirson, D.R. (1986). Histochemistry of lignin and suberin deposition in boundary layers formed after wounding in various plant species and organs. Can. J. Bot. 65: 1886–1892.

Robert, J.D. and Caserio M.C. (1977). Basic Principles of Organic Chemistry, second edition. W. A. Benjamin, Inc., Menlo Park, CA. ISBN 0-8053-8329-8.

Robb, J., Lee, S.W., Mohan, R., and Kolattukudy, P.E. (1991). Chemical characterization of stress-induced vascular coating in tomato. Plant Physiol. 97: 528–536.

Sabella, E., Luvisi, A., Aprile, A., Negro, C., Vergine, M., Nicolì, F., Miceli, A., and De Bellis, L. (2018). Xylella fastidiosa induces differential expression of lignification related-genes and lignin accumulation in tolerant olive trees cv. Leccino. J. Plant Physiol. 220: 60–68.

Salas-González, I., Reyt, G., Flis, P., Custódio, V., Gopaulchan, D., Bakhoum, N., Dew, T.P., Suresh, K., Franke, R.B., Dangl, J.L., Salt, D.E., and Castrillo, G. (2021). Coordination between microbiota and root endodermis supports plant mineral nutrient homeostasis. Science. 371: eabd0695.

Schmidt, A., Grimm, R., Schmidt, J., Scheel, D., and Strackt, D. (1999). Cloning and expression of a potato cDNAencoding hydroxycinnamoyl-CoA:tyramine N-(hydroxycinnamoyl)transferase. J. Biol. Chem. 274: 4273–4280.

Scortichini, M. (2020). The multi-millennial olive agroecosystem of salento (Apulia, Italy) threatened by Xylella fastidiosa subsp. Pauca: Aworking possibility of restoration. Sustain. 12: 6700.

Serra, O., Figueras, M., Franke, R., Prat, S., and Molinas, M. (2010). Unraveling ferulate role in suberin and peridermbiology by reverse genetics. Plant Signal. Behav. 5: 953–958.

Serrano, M., Coluccia, F., Torres, M., L’Haridon, F., and Métraux, J.P. (2014). The cuticle and plant defense to pathogens. Front. Plant Sci. 5: 274.

Street, P.F.S., Robb, J., and Ellis, B.E. (1986). Secretion of vascular coating components by xylem parenchyma cells of tomatoes infected with Verticilliumalbo-atrum. Protoplasma 132: 1–11.

Thoquet, P., Olivier, J., Sperisen, C., Rogowsky, P., Laterrot, H., and Grimsley, N. (1996). Quantitative trait loci determining resistance to bacterial wilt in tomato cultivar Hawaii7996. Mol. Plant-Microbe Interact. 9: 826–836.

Türker-Kaya, S. and Huck, C.W. (2017). Areview of mid-infrared and near-infrared imaging: principles, concepts and applications in plant tissue analysis. Molecules 22: 168.

Underwood, W. (2012). The plant cell wall: a dynamic barrier against pathogen invasion. Front. Plant Sci. 3: 85.

Ursache, R. et al. (2021). GDSL-domain proteins have key roles in suberin polymerization and degradation. Nat. Plants 7: 353–364.

VanderMolen, G.E., Beckman, C.H., and Rodehorst, E. (1987). The ultrastructure of tylose formation in resistant banana following inoculation with Fusariumoxysporum f.sp. cubense. Physiol. Mol. Plant Pathol. 31: 185–200.

Vasse, J., Frey, P., and Trigalet, A. (1995). Microscopic studies of intercellular infection and protoxylem invasion of tomato roots by Pseudomonas solanacearum. Mol. Plant-Microbe Interact. 8: 241–251.

Wang, J.F., Ho, F.I., Truong, H.T.H., Huang, S.M., Balatero, C.H., Dittapongpitch, V., and Hidayati, N. (2013). Identification of major QTLs associated with stable resistance of tomato cultivar “Hawaii 7996” to Ralstonia solanacearum. Euphytica 190: 241–252.

Wang, J.F., Olivier, J., Thoquet, P., Mangin, B., Sauviac, L., and Grimsley, N.H. (2000). Resistance of tomato line Hawaii7996 to Ralstonia solanacearum Pss4 in Taiwan is controlled mainly by a major strain-specific locus. Mol. Plant-Microbe Interact. 13: 6–13.

Xu, L., Zhu, L., Tu, L., Liu, L., Yuan, D., Jin, L., Long, L., and Zhang, X. (2011). Lignin metabolism has a central role in the resistance of cotton to the wilt fungus Verticilliumdahliae as revealed by RNA-seq-dependent transcriptional analysis and histochemistry. J. Exp. Bot. 62: 5607–5621.

Yadeta, K.A. and Thomma, B.P.H.J. (2013). The xylem as battleground for plant hosts and vascular wilt pathogens. Front. Plant Sci. 4: 97.

Zeiss, D.R., Piater, L.A., and Dubery, I.A. (2021). Hydroxycinnamate amides: intriguing conjugates of plant protective metabolites. Trends Plant Sci. 26: 184–195.

Zhang, Y., Zhang, W., Han, L., Li, J., Shi, X., Hikichi, Y., and Ohnishi, K. (2019). Involvement of a PadR regulator PrhP on virulence of Ralstonia solanacearumby controlling detoxification of phenolic acids and type III secretion system. Mol. Plant Pathol. 20: 1477–1490.

